# Loss of *IRF5* increases ribosome biogenesis leading to alterations in mammary gland architecture and metastasis

**DOI:** 10.1101/2023.05.01.538998

**Authors:** Zarina Brune, Dan Li, Su Song, Dan Iris Li, Isabel Castro, Rhea Rasquinha, Matthew R. Rice, Qin Guo, Kyle Kampta, Matthew Moss, Morgan Lallo, Erica Pimenta, Carter Somerville, Margaret Lapan, Victoria Nelson, Camila O. dos Santos, Lionel Blanc, Kevin Pruitt, Betsy J. Barnes

**Affiliations:** Center for Autoimmune Musculoskeletal and Hematopoietic Disease, The Feinstein Institutes for Medical Research, Manhasset, NY 11030; Donald and Barbara Zucker School of Medicine at Hofstra/Northwell, Hempstead, NY 11549; Department of Immunology and Molecular Microbiology, Texas Tech University Health Sciences Center, Lubbock, TX 79430; Department of Medical Oncology, Dana-Farber Cancer Institute, Boston, MA 02215; Broad Institute of MIT and Harvard, Cambridge, MA 02142; Cold Spring Harbor Laboratory, Cold Spring Harbor, NY 11724; Departments of Pediatrics and Molecular Medicine, Donald and Barbara Zucker School of Medicine at Hofstra/Northwell, Hempstead, NY 11549

**Author notes:** Equal contribution (Zarina Brune, Dan Li).

## Abstract

Despite the progress made in identifying cellular factors and mechanisms that predict progression and metastasis, breast cancer remains the second leading cause of death for women in the US. Using The Cancer Genome Atlas and mouse models of spontaneous and invasive mammary tumorigenesis, we identified that loss of function of interferon regulatory factor 5 (IRF5) is a predictor of metastasis and survival. Histologic analysis of *Irf5^-/-^* mammary glands revealed expansion of luminal and myoepithelial cells, loss of organized glandular structure, and altered terminal end budding and migration. RNA-seq and ChIP-seq analyses of primary mammary epithelial cells from *Irf5^+/+^* and *Irf5^-/-^*littermate mice revealed IRF5-mediated transcriptional regulation of proteins involved in ribosomal biogenesis. Using an invasive model of breast cancer lacking *Irf5*, we demonstrate that IRF5 re-expression inhibits tumor growth and metastasis via increased trafficking of tumor infiltrating lymphocytes and altered tumor cell protein synthesis. These findings uncover a new function for IRF5 in the regulation of mammary tumorigenesis and metastasis.

**Highlights:** Loss of IRF5 is a predictor of metastasis and survival in breast cancer.

IRF5 contributes to the regulation of ribosome biogenesis in mammary epithelial cells.

Loss of IRF5 function in mammary epithelial cells leads to increased protein translation.

## Introduction

The mammary gland is one of the most regenerative organs in the body, undergoing multiple transformations during the embryonic, pubertal, reproductive, and post-reproductive stages of life. Formation of the ductal tree, the functional backbone of the mammary gland, is orchestrated by a specialized structure called the terminal end bud (TEB). TEBs are bulb-shaped structures unique to mammary glands that direct the growth of ducts throughout the fat pad (Paine and Lewis, 2017). In addition, TEBs are responsible for the production of differentiated cells, including myoepithelial (basal) cells and luminal cells. These mature cell types support ductal elongation and are a regulatory point for basement membrane deposition, branching, angiogenesis, and pattern formation. The counterpart of mammary gland development is a process called involution. Described as the reverse of development, involution is a complex multistage process characterized by regression of the mammary gland epithelium to its non-lactating state through apoptosis and tissue remodeling. Mammary gland tissue remodeling during involution mimics pathological conditions of wound healing, inflammation, and tumorigenesis. Given the unique characteristics of TEBs, including their heterogenous cellular composition, high proliferation and apoptosis rates, invasive ability, and angiogenic properties, alterations in TEB function(s) have been linked to mammary tumorigenesis. Thus, although hormonal control of TEB growth is well-characterized, the factors that regulate ductal elongation and outgrowth require further attention and evaluation as potential drivers of mammary tumorigenesis and metastasis.

Breast cancer (BC) is the most common cancer in women worldwide. Since 2008, BC incidence has increased by more than 20% and mortality by 14% (DeSantis et al., 2015). In the U.S., 1 in 8 women will develop metastatic BC or invasive ductal carcinoma (IDC) (Siegel et al., 2022). This equates to an estimated 287,850 new cases of invasive disease diagnosed in 2022, along with 51,400 new cases of non-invasive BC (Siegel et al., 2022). Second to lung cancer, mortality from BC represents the highest rate of cancer death for women in the U.S. Although most new cases of BC are invasive, or infiltrating, ∼one-third or more of the non-invasive ductal carcinoma *in situ* (DCIS) cases will progress to IDC if left untreated (Siegel et al., 2022). Given that metastasis of primary BC to distant sites and the recurrence of therapeutically recalcitrant disease are the main causes of fatalities, a clear understanding of the molecular mechanisms controlling BC development and progression is essential. Equally important is the identification of prognostic markers that can delineate patients who are at the highest risk for developing metastasis and have the potential to respond to targeted therapy. Extensive characterization of dysregulated factors and pathways driving BC has been accomplished using modern sequencing technologies. Despite this, only ∼5-10% of BCs have been linked to inherited gene mutations. These data strongly suggest that a significant proportion of the mechanisms contributing to BC susceptibility and mortality are not associated with genomic mutation.

The transcription factor interferon regulatory factor 5 (IRF5) is a critical mediator of the immune response to pathogens, cellular response to DNA damage, and a tumor suppressor (Barnes et al., 2003; Barnes et al., 2001; Barnes et al., 2004; Bi et al., 2011; Brune et al., 2020; Cevik et al., 2017; Dai et al., 2017; Dong et al., 2015; Fresquet et al., 2012; Hu et al., 2005; Mori et al., 2002; Wies et al., 2009; Yanai et al., 2007). In the context of tumorigenesis, we previously reported that normal breast tissue and tissue with atypical ductal hyperplasia (ADH) stained positive for IRF5 expression, whereas only ∼38% of DCIS and ∼10% of IDC retained IRF5 expression (Bi et al., 2011; Pimenta et al., 2015). A correlation analysis of *IRF5* transcript expression with recurrence-free survival (RFS) in all BCs in The Cancer Genome Atlas (TCGA-BRCA) revealed that the lower quartile of *IRF5* expression was a significant prognostic marker of poor RFS (Pimenta and Barnes, 2015).

To further elucidate the role of IRF5 in BC, we examined spontaneous mammary tumorigenesis in female littermate-matched *Irf5^+/+^* and *Irf5^-/-^* BALB/c mice and utilized the 4T1 model of invasive BC to examine IRF5-mediated metastasis. From these studies, we found that IRF5 is involved in mammary gland development, BC initiation, metastatic colonization, and survival. We demonstrate a role for IRF5 in various stages of murine mammary gland development and maturation, including a role in pregnancy remodeling and involution. Furthermore, our findings demonstrate a correlation between the retention of IRF5 expression in basal or triple-negative BC (TNBC) and increased overall survival (OS) and RFS. Finally, we identify a previously undescribed function for IRF5 in the regulation of ribosome biogenesis and protein translation in both primary and immortalized mammary gland epithelium, providing a mechanistic link between IRF5 expression, tumorigenesis, and metastasis.

## Results

### IRF5 expression is a marker of OS and RFS in human breast cancer

Utilizing bulk RNA-seq data from matched normal breast epithelial and cancer cells from the TCGA-BRCA database, we reported a decrease in *IRF5* expression that correlated with decreased RFS (Pimenta and Barnes, 2015). To exclude potential confounders due to tumor-associated immune cells that may express *IRF5*, we refined our analysis utilizing publicly available scRNA-seq datasets of matched normal breast epithelial and cancer cells. As expected, we observed a significant reduction in tumor-intrinsic *IRF5* expression compared to normal breast epithelial cells (**Fig. 1A, B**); trends towards reduced *IRF5* expression in IDC compared to DCIS were also detected (**Supp. Fig. 1A**) (Bi et al., 2011; Pimenta et al., 2015).

**Figure 1:**
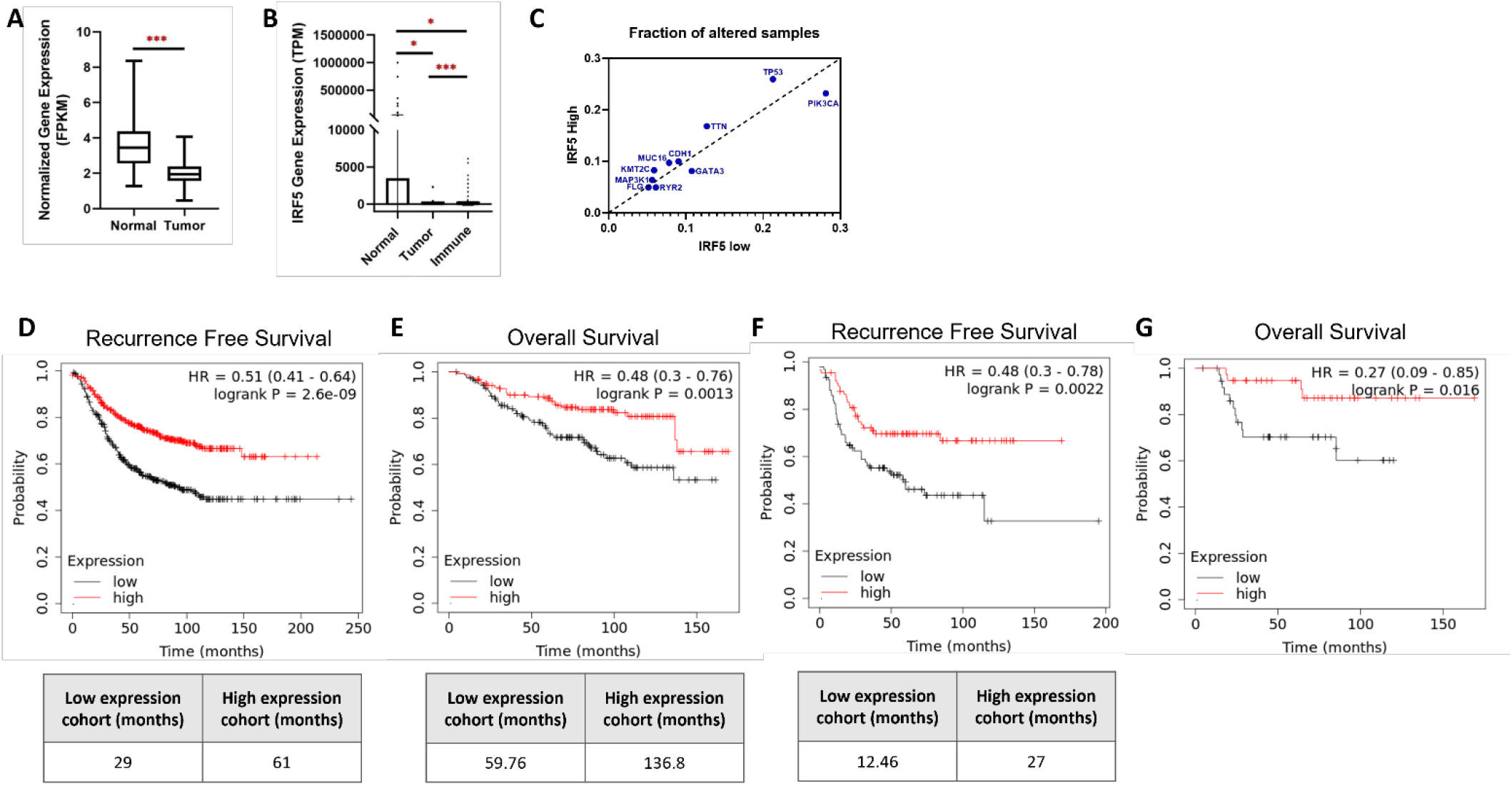
*IRF5* expression is a marker of OS and RFS in human breast cancer. **(A, B)** IRF5 expression is lost in breast cancer. **(A)** IRF5 expression was determined from bulk RNA-seq data using GDC TCGA-BRCA downloaded from the UCSC Xena portal (n=1097) and matched normal data GSE86354 (n=92). Boxplot representation of IRF5 expression normalized to FPKM. **(B)** IRF5 expression determined by cell type using published scRNA-Seq datasets (Normal (GSE113196), Tumor (GSE75688), and Immune (GSE114725)). Graphical representation (individual values with mean ± SD) of single-cell IRF5 expression data normalized to TPM in normal epithelial (*n* = 867), breast cancer tumor (*n* = 1,851) and tumor-associated immune cells (*n* = 21,253). \**p*< 0.05; \*\**p*< 0.01; \*\*\**p*< 0.0001. **(C)** IRF5 expression is not associated with common BC driver mutations. **(C)** Data are from TCGA-BRCA (Cell 2015, Firehose legacy, Nature 2012, and PanCancer Atlas) and METABRIC datasets. High, IRF5 expression ≥2 standard deviations (SD) above normal matched breast tissue; Low, IRF5 expression <2 SD below control samples. Distribution plot of BC mutations in the TCGA-BRCA cohort of patients with either low or high IRF5 expression. For PIK3CA: 0.28% of IRF5 low expressing tumors show mutations within this gene and 0.23% of the IRF5 high expressing tumors. For TP53: 0.21% of the IRF5 low group vs. 0.26% of the IRF5 high group. **(D-G)** Upper quartile of human *IRF5* tumoral expression is a significant marker of prolonged RFS and OS. **(D)** Data of RFS are from n=1764 patients within the TCGA-BRCA cohort. Black line is lower quartile of IRF5 expression, red line is upper quartile. **(E)** Same as **D** except data are for OS from n=626 patients. **(F, G)** Similar to **(D, E)** except data of RFS are from n=360 **(F)** and OS are from n=153 **(G)** patients with TNBC from TCGA-BRCA and METABRIC. Graphs are from the Kaplan-Meier Database; JetSet best probe set used (IRF5 239412_at). Tables under the survival curves **(D-F)** are the upper quartile survival (months) for the low expression (bottom 25%) and high expression (top 25%) groups.

We next examined whether reduced expression was due to putative copy-number alterations, loss of heterozygosity, or mutation(s) in the *IRF5* gene. Utilizing TCGA-Breast Invasive Carcinoma (TCGA-BIC) cohort, we determined that 94% of IDC were *IRF5* low and ≤1% of IDCs had a genetic modification (**Table I**). These data suggest that decreased *IRF5* expression in IDCs is likely due to epigenetic modification(s) and/or chromatin remodeling (Li and Tainsky, 2011; Li et al., 2008; Schlosser et al., 2021; Yamashita et al., 2010). Furthermore, we found that *IRF5* expression patterns in BC were not correlated with classical BC driver mutations, including *MYC*, *CCND1*, *HER2* (*ERBB2*), *TP53*, *PIK3CA*, and *BRCA1/2* (**Fig. 1C**), suggesting that loss of *IRF5* expression in IDC is an independent event in BC tumorigenesis.

**Table I.** Characterization of IRF5 expression in mammary tumors.

| **IRF5 expression<sup>2</sup>** | **% of tumors** | **Mutation** | **% of TILS<sup>3</sup>** |
| --- | --- | --- | --- |
| **High (IRF5-positive)** | **6%** | **0** | **58%** |
| **Low (IRF5-negative)** | **94%** | **≤1%** | **22%** |

<sup>1</sup> Data are from The Cancer Genome Atlas-Breast Invasive Carcinoma group, Cell 2015 (n=1105). <sup>2</sup>High, IRF5 expression $\geq 2$ standard deviations (SD) above normal matched breast tissue; Low, IRF5 expression $< 2$ SD below control samples. <sup>3</sup>TIL determination based on individual patient pathology reports. Shown are % of tumors with TILs.

Studies of gene expression analysis in human BC identified *IRF5* within a molecular signature that predicted increased survival and lower incidence of metastasis (Ma et al., 2007; van ’t Veer et al., 2002). We next utilized the TCGA-BRCA cohort to examine the correlation between *IRF5* expression and patient survival in different BC subtypes, including hormone-receptor positive – Luminal A (estrogen-receptor positive (ER+), progesterone-receptor positive (PR+), HER2-negative (HER2-), Ki-67-low) and Luminal B (ER+, PR+, HER2+/-, Ki-67-high), hormone-receptor negative – Basal-like (ER-, PR-, HER2-; TNBC) and HER2-enriched (ER-, PR-, HER2+). Kaplan-Meier survival curves of BC patients were generated comparing high and low *IRF5* expression (top and bottom 25%, respectively). We found that those patients with the top 25% of *IRF5* expression had significantly increased RFS (*p*=2.6×10^-9^) and OS (*p*=0.0013) (**Fig. 1D, E**). Similarly, when we stratified BCs by subtype, we found that the upper *IRF5* expression quartile was a significant marker of prolonged RFS (*p*=0.0022) and OS (*p*=0.016) in TNBC (**Fig. 1F, G**), providing a 2-fold increase in life expectancy for patients within this cohort. Similar findings were made in the other subtypes for RFS but not OS (**Supp. Fig. 1B-G**). Together, these data provide valuable predictive insights into disease course for *IRF5* in TNBC, a cancer that represents 15-20% of all BCs, is highly metastatic, has worse clinical outcomes, and remains challenged by limited therapeutic options (Dent et al., 2007; Lehmann et al., 2011).

### Aging nulliparous Irf5^-/-^ Balb/c mice have increased incidence of spontaneous mammary tumorigenesis

To examine the mechanistic link between *IRF5* expression and BC development, we generated a colony of *Irf5^+/+^* (WT) and *Irf5^-/-^* (KO) littermate-matched Balb/c mice. Female 12 months-old WT Balb/c mice have a reported 1-4% incidence of spontaneous mammary tumorigenesis that increases to >7% in response to low-dose whole-body γ-irradiation (Medina, 2010; Yu et al., 2001). Others and we previously reported p53-dependent and -independent tumor suppressor roles for IRF5 in DNA damage-induced apoptosis (Barnes et al., 2003; Bi et al., 2014; Bi et al., 2011; Cevik et al., 2017; Fresquet et al., 2012; Hu and Barnes, 2009; Hu et al., 2005; Mori et al., 2002; Yanai et al., 2007). To confirm that loss of *Irf5* leads to increased DNA damage-induced tumorigenesis, 12 weeks-old nulliparous WT and KO Balb/c mice underwent whole-body γ-irradiation and were euthanized at 12 months of age for histologic mammary gland analysis. We detected the expected rate of 4% (n=25) basal spontaneous mammary tumorigenesis in mock-irradiated WT mice (**Table II**) that increased in incidence to 11% (n=9) following γ-irradiation (**Suppl. Table I**). Conversely, KO mice had a 14% (n=21) incidence of spontaneous mammary tumorigenesis that increased to 43% (n=7) following γ-irradiation (**Table II**, **Suppl. Table I**). These data support a critical protective role for *Irf5* in both spontaneous and DNA damage-induced mammary tumorigenesis.

**Table II.**
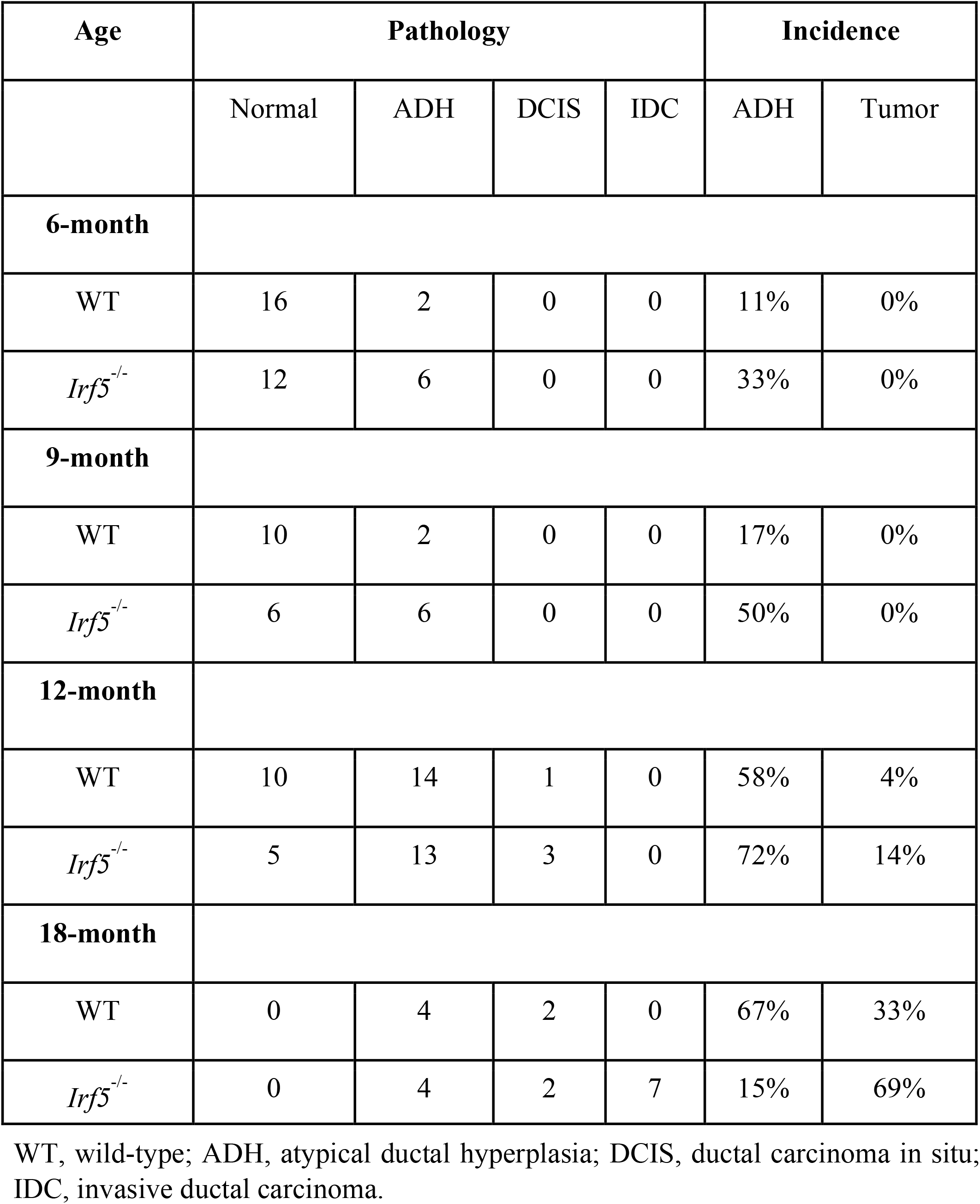
Spontaneous tumor incidence in female littermate matched WT and *Irf*5^-/-^ Balb/c mice.

Given that *IRF5* expression associates with the clinical prognosis of BC (**Fig. 1**), we performed an age-dependent analysis of mammary tumorigenesis in female nulliparous littermate-matched WT and KO mice. We examined mammary tumor formation in 6, 9, 12, and 18 months-old mice by hematoxylin and eosin (H&E), immunohistochemistry (IHC) and immunofluorescence (IF) analysis. At 6 months-old, KO mice had a 33% incidence of spontaneous ADH (n=18); a rate three-fold higher than that seen in WT littermates (n=18). Spontaneous ADH incidence further increased to 17 and 50% in 9 months-old WT (n=12) and KO (n=12) mice, respectively. Strikingly, 18 months-old WT mice (n=6) had a 67% incidence of spontaneous ADH and a 33% tumor incidence, while KO mice (n=13) had a 15% incidence of ADH and 69% incidence of tumor formation (**Table II**, **Fig. 2A**). These data indicate that with increasing age, loss of *Irf5* on a Balb/c background leads to a significant increase in the rate of spontaneous mammary tumorigenesis. Characterization of hormone receptor status in histologically normal mammary glands from 18 months-old WT and KO mice revealed similar ER+, PR+ and HER2-low staining while tumors isolated from 18 months-old KO mice were ER-low, PR-, HER2- (**Fig. 2B, C**). IF analysis showed expansion of KO luminal epithelial cells with increased Ki67 staining, suggesting luminal cell origin for the KO primary tumors (**Fig. 2D-F**, **Supp. Fig. 2**).

**Figure 2:**
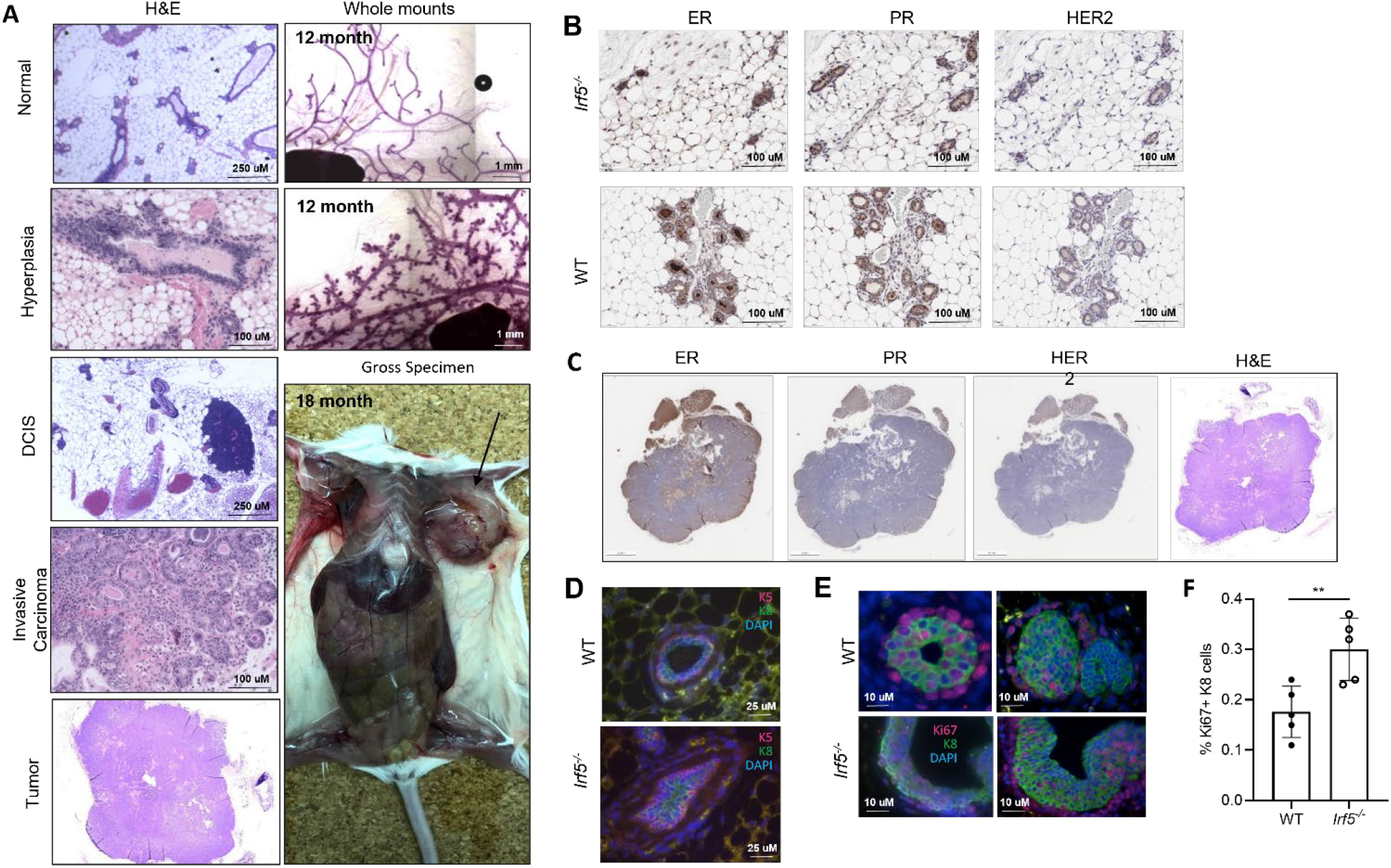
*Irf5^-/-^* Balb/c mice have increased incidence of spontaneous mammary tumorigenesis. **(A)** Representative H&E, whole mount, and gross specimen of mammary glands from WT and *Irf5^-/-^* mice at the indicated age and/or tumor stage; n=4/genotype. Gross spontaneous tumor is shown in 18 months-old *Irf5^-/-^*mouse (black arrow). **(B, C)** Representative IHC staining of mammary glands for hormone receptor expression in 18-month-old healthy WT and *Irf5^-/^*^-^ mammary glands **(B)** and tumor from *Irf5^-/-^* mammary gland **(C)**; n=4/genotype. (ER = Estrogen receptor, PR = Progesterone receptor, HER2 = Human epidermal growth factor receptor 2). **(D)** Representative immunofluorescence (IF) staining of 12-month WT and *Irf5^-/-^* mammary glands (n=4/genotype); K5 (basal; red) and K8 (luminal; green) demonstrating expansion of both compartments. **(E)** Representative IF staining of 12-month WT and *Irf5^-/-^* mammary glands (n=5/genotype); Ki67 (red), K8 (luminal; green) and DAPI (blue) demonstrating increased luminal epithelial cell proliferation. **(F)** Quantification of Ki67 expression from **(E)**; n=5/genotype. \*\**p*< 0.001.

### Loss of Irf5 expression alters normal mammary gland development and architecture

IRF5 has been characterized as a key transcriptional mediator of type I interferons (IFNs) and other pro-inflammatory cytokines downstream of Toll-, NOD-, and RIG-I-like receptors. In addition, in autoimmune diseases such as systemic lupus erythematosus (SLE), inflammatory bowel disease (IBD) and rheumatoid arthritis (RA), increased genetic risk is associated with increased IRF5 expression and hyper-activation in circulating immune cells (Byrne et al., 2017; Fabie et al., 2018; Feng et al., 2012; Krausgruber et al., 2011; Li et al., 2020; Song et al., 2020; Weiss et al., 2015; Yan et al., 2020). To begin to understand how loss of *Irf5* contributes to murine mammary tumorigenesis, we examined IRF5 expression in WT mammary glands from 6 weeks-old mice. Using RNA fluorescent *in situ* hybridization (RNA-FISH), we show that *Irf5* mRNA is expressed in both myoepithelial and luminal cells, albeit at increased levels in myoepithelial cells (**Fig. 3A**, **Suppl. Fig. 3A**). These findings were confirmed by qPCR on sorted cell populations (**Fig. 3B**, **Suppl. Fig. 3B**). We next examined IRF5 protein expression by IHC, Western blot and imaging flow cytometry analysis. Like *Irf5* transcript expression, IRF5 protein levels were increased in myoepithelial cells compared to luminal cells, revealing primarily cytoplasmic expression (**Fig. 3C-G**).

**Figure 3:**
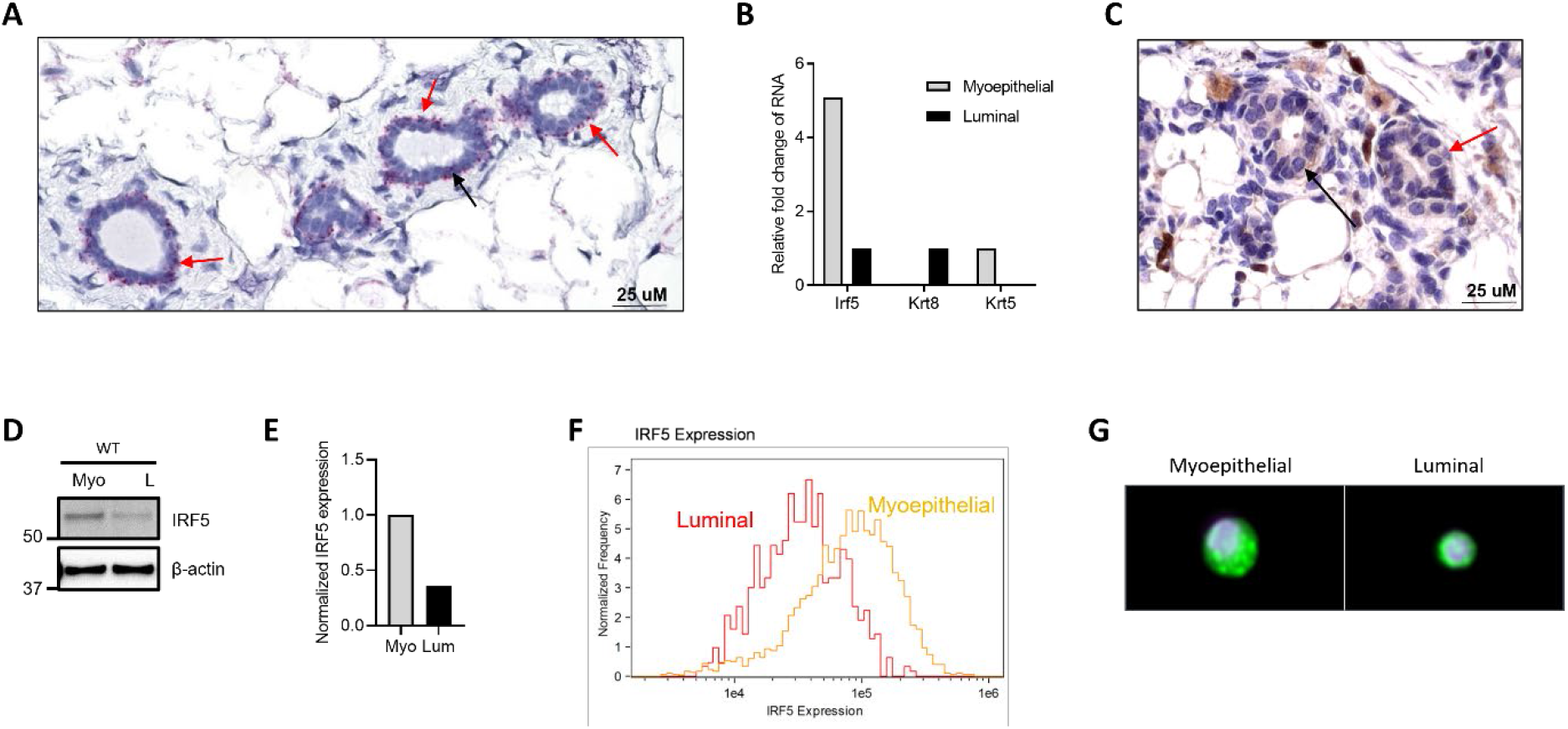
IRF5 is expressed in both myoepithelial and luminal cells. **(A)** Representative *Irf5* RNA *in situ* hybridization of WT mammary glands at 40X magnification. *Irf5* is more highly expressed in myoepithelial cells (red arrow); luminal epithelial cells (black arrow). n=3 independent replicates. **(B)** Representative *Irf5* qRT-PCR from sorted WT myoepithelial and luminal cells. Data are representative of n=3 independent replicates. **(C)** Representative IHC staining of IRF5 in WT mammary glands. Myoepithelial (red arrow) and luminal (black arrow) cells are shown; n=3 independent replicates. **(D, E)** Representative Western blot **(D)** and associated quantification **(E)** of IRF5 expression in sorted myoepithelial (Myo) and luminal (L) cells pooled from n=4 WT mice. n=2 independent replicates. **(F, G)** Representative histogram of IRF5 expression **(F)** and cellular localization **(G)** in myoepithelial and luminal cells as determined by imaging flow cytometry. Images captured at 60X magnification. IRF5 is shown in green and nucleus by DAPI (blue). ≥500 captured cells from 2 independent replicates.

We next examined the role of IRF5 in normal mammary gland development. Whole mount preparations and histology from age-matched WT and KO mammary glands showed increased fat pad filling with increased ductal branching, TEB numbers, and increased TEB migration beginning at the pre-pubertal age of 4.5-weeks (**Fig. 4A, B**). Increased TEB numbers and secondary branching occurred in KO mice throughout all stages of mammary gland development (**Fig. 4A-C**). To assess whether a global loss of *Irf5* altered the immune microenvironment of aging mammary glands, we immunophenotyped WT and KO mammary glands from 9 months-old mice using flow cytometry. Although we found no significant differences in immune cell populations between WT and KO mammary glands, after sub-setting macrophages based on markers of M1 and M2 polarization, we detected a significant reduction in M1 macrophages from KO mammary glands (**Supp. Fig. 4**). To clearly show that alterations in TEB formation and migration were due to an intrinsic loss of *Irf5* in mammary epithelial cells, mammary duct organoids from 6 weeks- and 9 months-old WT and KO mice were generated. For both ages, we found a significant increase in organoid branching from KO mice compared to their WT counterpart (**Fig. 5A-D**). Together, these data support an intrinsic role for IRF5 in TEB formation and migration.

**Figure 4:**
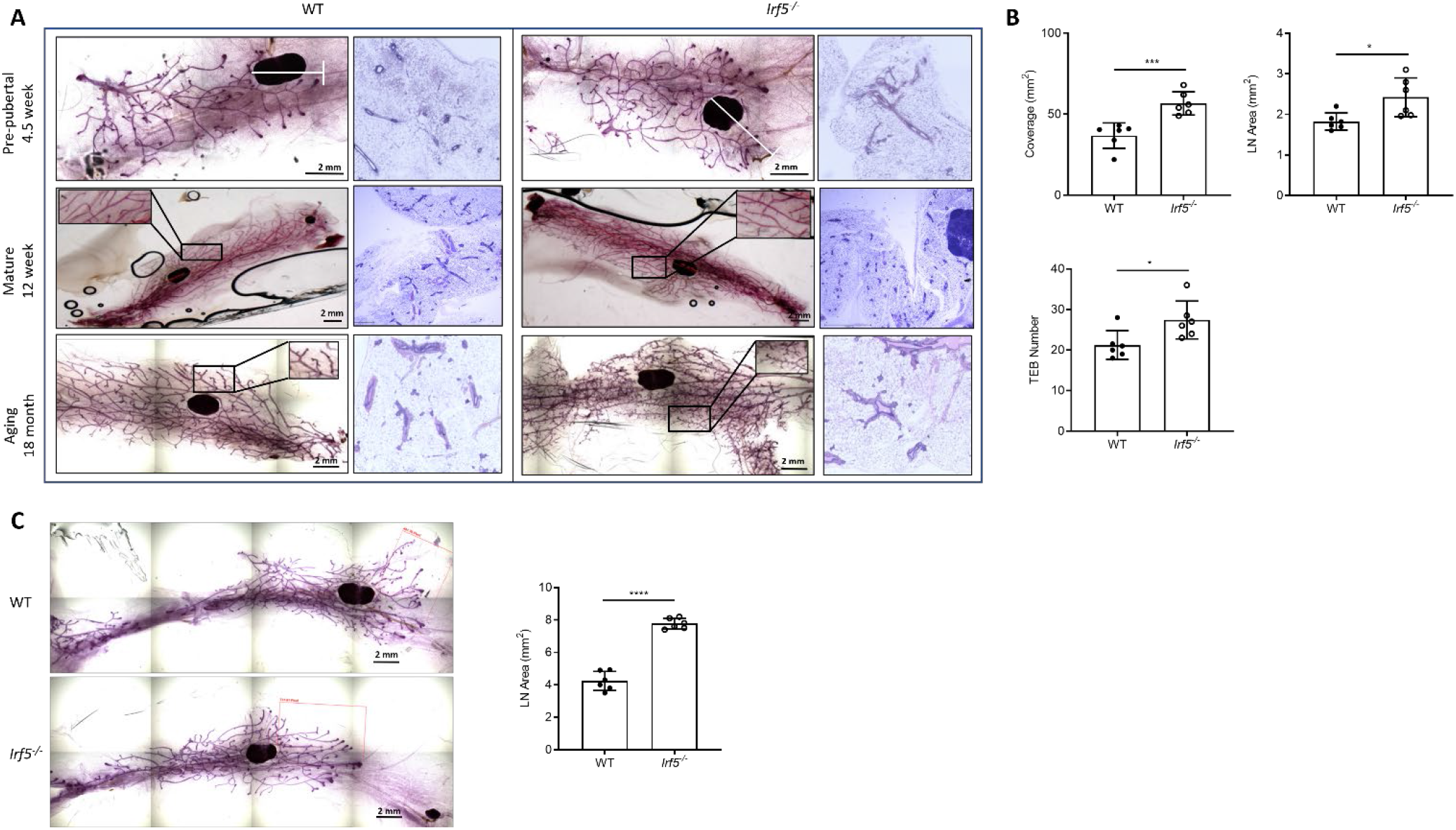
Loss of *Irf5* expression alters normal mammary gland structure during aging. **(A)** Representative whole mount with H&E staining of WT and *Irf5^-/-^* mammary glands at indicated ages. Scale bar for whole mounts represents 2 mm; n = 6 mice/age/genotype. **(B)** Quantification of fat pad filling by measuring total coverage (Coverage), terminal end bud (TEB) migration from lymph node (LN) area and absolute TEB numbers in 4.5 weeks-old pre-pubertal mice are shown. \**p*< 0.05, \*\**p*< 0.01; n = 6 mice/genotype. **(C)** Same as **(A)** except representative whole mounts from 6 weeks-old mammary glands are shown with quantification of terminal end bud (TEB) migration relative to the LN. Scale bar represents 2 mm. \*\*\*\**p*< 0.0001. (n = 6 mice/genotype)

**Figure 5:**
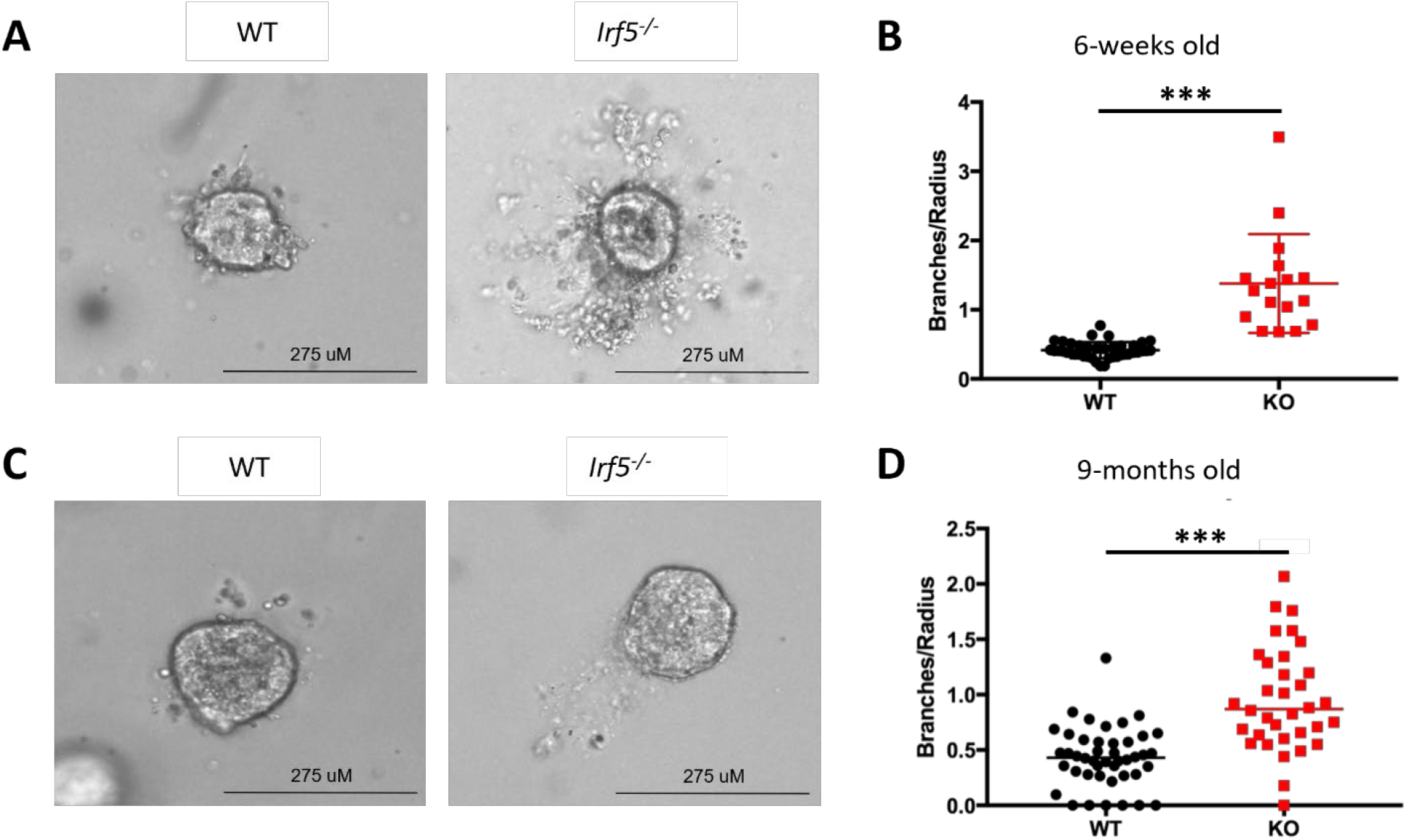
Loss of *Irf5* expression results in increased budding and invasiveness of *Irf5^-/-^* mammary organoids. **(A – D)** Representative images and quantification of mammary epithelial cell budding/branching within *in vitro* organoid cultures from mammary glands of 6 weeks-old **(A, B)** and 9 months-old **(C, D)** WT and *Irf5^-/-^* Balb/c mice. Pictures were taken at Day 4 of growth. n = 3/genotype performed in triplicate. \*\*\**p*< 0.0001.

Lastly, flow cytometry analysis was performed to directly characterize the myoepithelial and luminal cell populations from WT and KO mammary glands. We first focused on luminal epithelial cells; the cell population responsible for the bulbous TEB structure (Visvader and Stingl, 2014). Similar to data in **Fig. 2D-F**, we detected a significant increase in total CD24^hi^CD29^lo^ luminal cells from the Lin^-^ population of 6 weeks-old KO mammary glands (**Suppl. Fig. 5A, B**). The number of Lin^-^CD24^hi^CD29^lo^CD61^+^CD133^-^ luminal progenitor cells was decreased. Furthermore, there were significantly more Lin^-^ CD24^hi^CD29^lo^CD61^-^CD133^+^ luminal ductal hormone responsive cells in KO compared to WT mammary glands (**Suppl. Fig. 5C**) (dos Santos et al., 2013). Interestingly, analysis of myoepithelial cell populations revealed a trend towards reduced KO mammary stem cells (MaSCs; Lin^-^CD24^+^CD29^hi^CD61^low^Cd1d^+^), a significant reduction in differentiated myoepithelial cells (Lin^-^CD24^+^CD29^hi^CD61^-^CD1d^-^) and increased myoepithelial progenitor cells (Lin^-^CD24^+^CD29^hi^CD61^+^Cd1d^-^) (**Supp. Fig. 5D**). These results support a role for IRF5 in the regulation of luminal and myoepithelial cell stemness, maturation, and TEB development.

Pregnancy has been shown to both promote and protect against BC (Slepicka et al., 2019). Studies have shown that a first full-term birth before the age of 20 or 25 can reduce the long-term risk of breast cancer by 50% or 38%, respectively (Katz, 2016). However, this protection does not occur until approximately 3-8 years following the last pregnancy (Breast Cancer Risk After Recent Childbirth, 2019). Multiple animal models of mammary tumorigenesis and pregnancy have confirmed the protective effects of pregnancy in BC (Feigman et al., 2020; Hanasoge Somasundara et al., 2021). Interestingly, prolonged lactation and nursing have also been identified as key risk factors for inflammatory breast cancer (IBC), a rare and highly aggressive BC subtype (Mejri et al., 2020). On the other hand, studies have shown that short-term breast feeding and premature involution may also contribute to a “primed” stroma that is permissive to IBC (Bambhroliya et al., 2018). During the process of post-weaning involution, mammary epithelial cells undergo programmed apoptosis and clearance by macrophages and phagocytic epithelial cells (Monks and Henson, 2009). Due to the observed alterations in development and tumorigenesis in KO mammary glands, as well as the known role for IRF5 in inflammation and macrophage polarization, we next examined if loss of *Irf5* impacted mammary gland remodeling and tumorigenesis post-pregnancy.

To examine the role of IRF5 in pregnancy driven mammary gland remodeling, a cohort of 3 months-old WT and KO breeder females underwent two timed pregnancies and then were aged to 12 months. Histologic analysis of mammary glands revealed that 0% of WT (n=7) compared to 37.5% of KO (n=8) breeder mice had mammary tumors (**Table III**). To further analyze the role of IRF5 in pregnancy-induced mammary gland remodeling, we performed histologic analysis of whole mount preparations of mammary glands from pregnant and involuting mice. KO mice had increased budding of the terminal alveoli at day five of pregnancy, with no significant gross structural alterations at day 15 of pregnancy (**Fig. 6A**). Following forced involution induction by early weaning, KO mice demonstrated few differences in mammary gland structure at day 5 post-induction, while at day 10 post-induction, residual branching and increased TEBs remained in KO mammary glands as compared to matched WT mammary glands, supporting defective or delayed involution in KO mice (**Fig. 6B**). Histologic analysis of involuting mammary glands indicated maintained dilation of mammary gland lumens with retention of milk production in KO mice at both days 5 and 10 compared to corresponding WT mouse mammary glands (**Fig. 6C**, **Supp. Fig. 6A**). To determine if alterations in immune cell infiltration were responsible for alterations in involution induced mammary gland remodeling, IHC was used to analyze CD45^+^ and F4/80^+^ infiltration at these same timepoints. There were no significant differences in immune cell infiltration into the involuting mammary glands at either day 5 or day 10 of involution (**Supp. Fig. 6B, C**).

**Figure 6:**
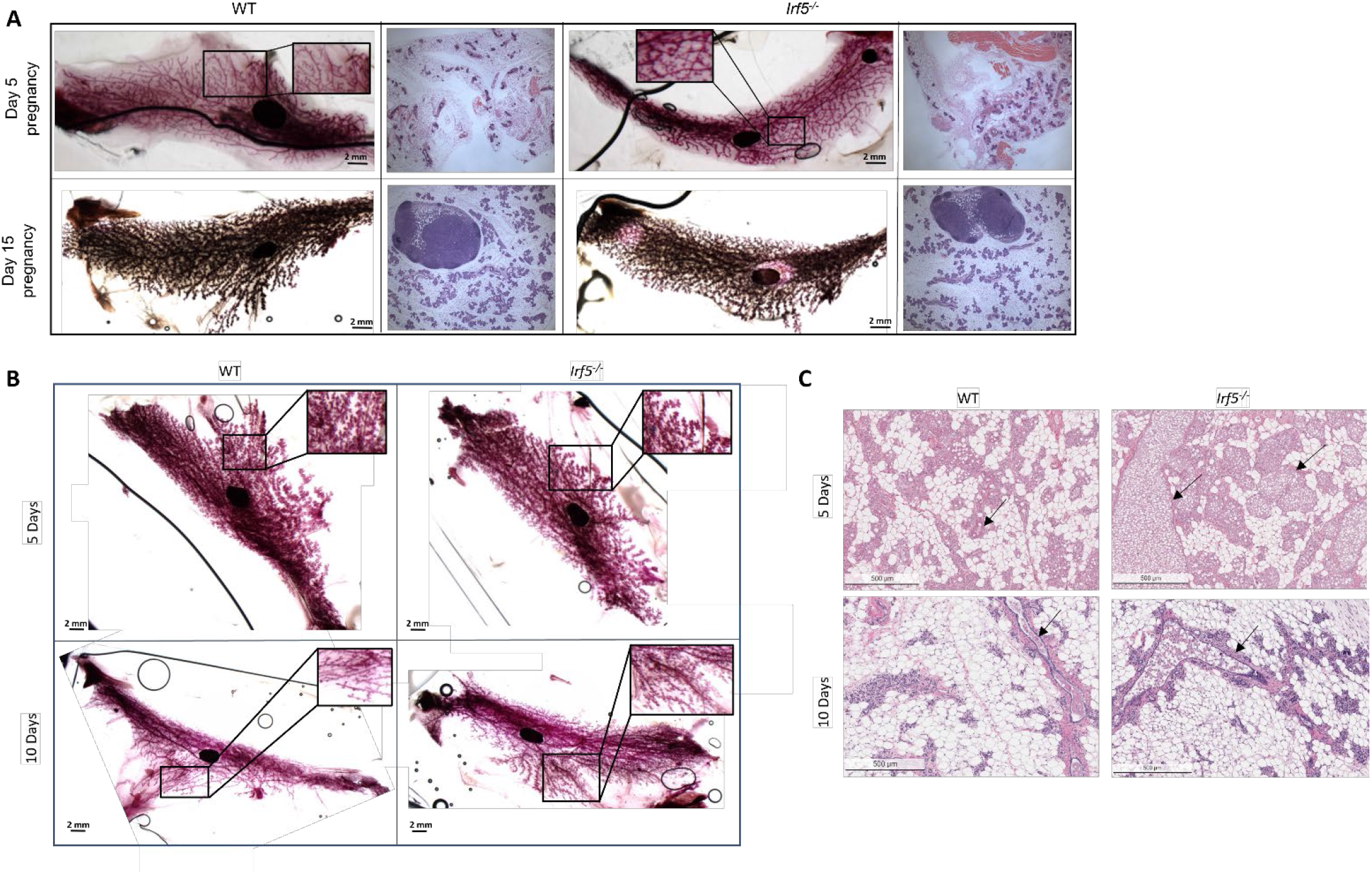
Alterations in mammary gland structure during pregnancy and involution contributes to increased incidence of mammary tumorigenesis in *Irf5^-/-^* mice. **(A)** Representative whole mounts of WT and *Irf5^-/-^* mammary glands with H&E staining at Days 5 and 15 of pregnancy. Scale bar for whole mounts represent 2 mm; n=3/time point/genotype. **(B)** Similar to **(A)** except whole mounts are showing mammary glands after forced involution induction at Days 5 and 10 post-weaning. Scale bar represents 2 mm; n=3/time point/genotype. **(C)** Representative H&E staining of involuting mammary glands from WT and *Irf5^-/-^* mice at Days 5 and 10 post-weaning revealing maintained dilation of *Irf5^-/-^* mammary gland lumens. Scale bar represents 500 µM; n = 3 mice/genotype.

**Table III.**
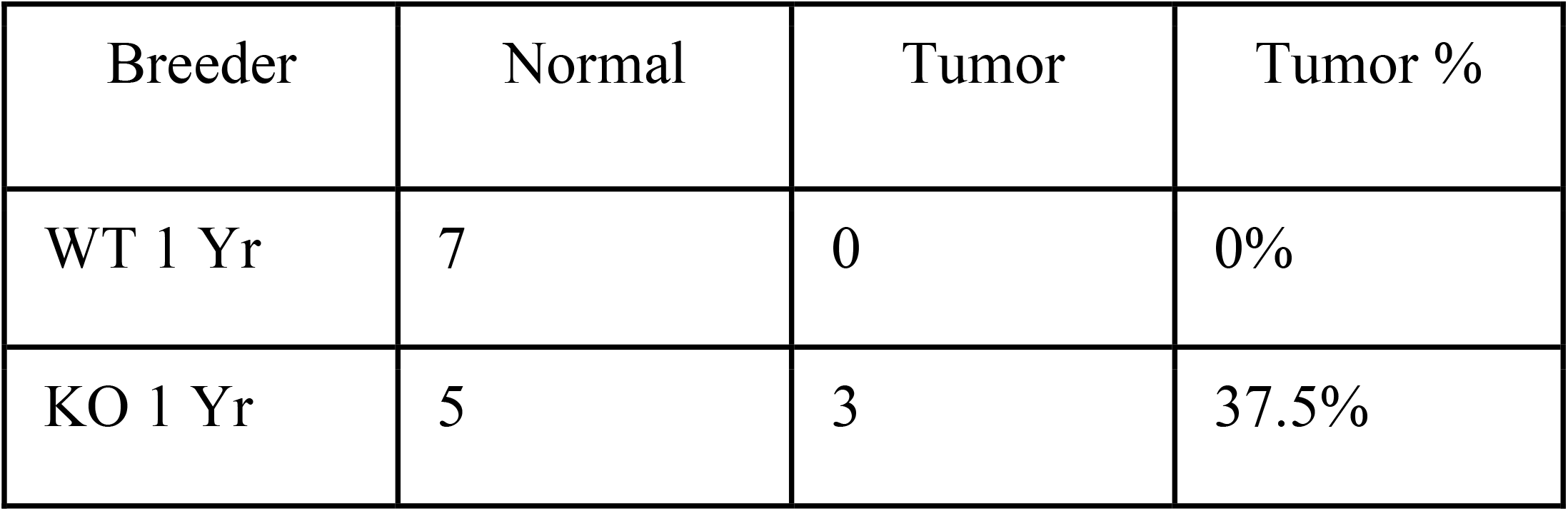
Spontaneous tumor incidence in breeder female littermate matched WT and *Irf*5^-/-^ Balb/c mice.

The process of involution is a coordinated program of cell death, mammary gland remodeling and immune cell infiltration (Wallace et al., 2019). We next sought to confirm that the involution defects corresponded with intrinsic defects in mammary epithelial cell programming upon loss of *Irf5*, rather than due to alterations in IRF5-mediated immune cell function. Organoids generated from 10 weeks-old WT and KO littermate mice were differentiated in a lactation inducing environment for five days as previously described (Sumbal et al., 2020). At day 5 following involution induction, organoids were harvested, stained, and analyzed using confocal microscopy. Analysis revealed KO organoids to have a dilated morphology compared to WT, further supporting an intrinsic defect in the ability of KO mammary epithelial cells to undergo normal involution processes (**Supp. Fig. 6D**).

### Intratumoral IRF5 expression inhibits in vivo tumor growth and metastasis

Given the significant correlation between IRF5-low BC and metastatic disease and the dysregulated growth and development of KO mammary glands, we next interrogated if loss of *Irf5* could drive primary tumor growth and metastasis *in vivo*. We utilized the orthotopic 4T1 model of metastatic BC that represents a stage IV TNBC. As expected, endogenous IRF5 expression in 4T1 cells was nearly undetectable, mimicking an IRF5-low TNBC (**Suppl. Fig. 7A**). We next generated stable IRF5-expressing 4T1 (IRF5-high) and 4T1.2-Luc3 cell lines for functional comparison to 4T1 IRF5-low cell lines; the level of stable overexpression was similar to expression in WT primary mammary epithelial cells (**Fig. 3D**). We first performed an *in vitro* characterization of 4T1 cell lines and found no difference in cell viability or proliferation between IRF5-high and -low cells (**Fig. 7A, B**). However, transwell assays showed a significant increase in migration and invasion of 4T1 IRF5-low cells compared to 4T1 IRF5-high cells (**Fig. 7C, D**). These data are reminiscent of findings in human breast cancer cell lines with low or high IRF5 expression predicting increased or decreased metastasis, respectively (Bi et al., 2011; Pimenta et al., 2015). We next orthotopically implanted 0.1×10^6^ IRF5-high or -low 4T1.2-Luc3 cells into mammary fat pads of WT female Balb/c mice and monitored tumor formation and lung metastasis by live bioluminescence imaging (BLI). Weekly monitoring revealed differences in both primary tumor growth and lung metastasis (**Fig. 7E**, **Supp. Fig. 7B**). Similar data were obtained with parallel experiments performed in IRF5-high and -low expressing 4T1 cell lines. At day 30 post-implantation, we detected significant reductions in tumor weight and lung metastatic nodules in 4T1 IRF5-high implanted mice (**Fig. 7F-H**). Further, we observed a significant increase in the survival of mice implanted with 4T1 IRF5-high tumors following primary tumor resection at day 10 post-implantation, compared to mice implanted with IRF5-low tumors (**Fig. 7I**). Altogether, these data indicate that low IRF5 tumor expression is both a significant risk factor for metastasis and a negative predictor of survival.

**Figure 7:**
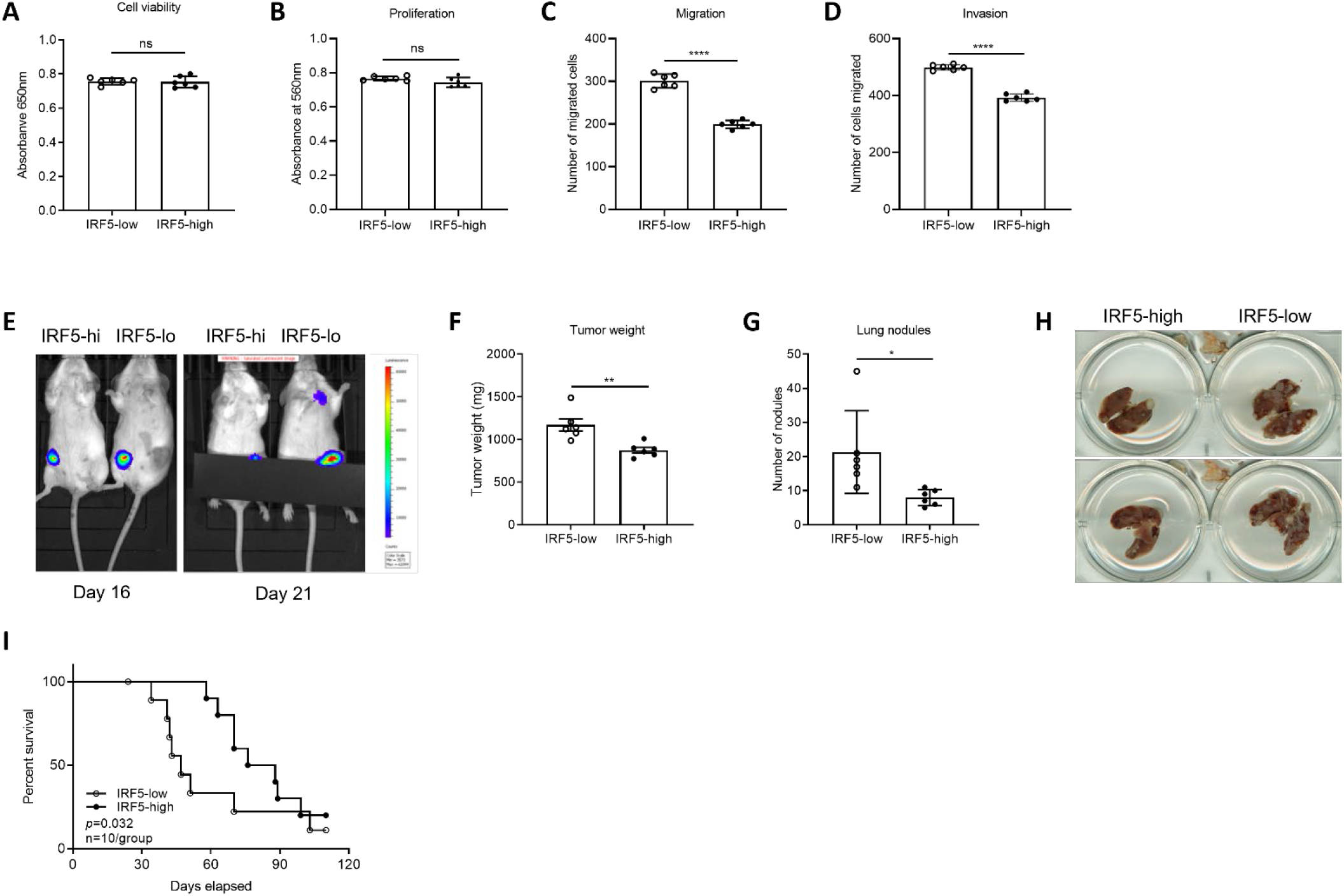
Intratumoral IRF5 expression inhibits *in vivo* tumor growth and metastasis. **(A)** Results from MTT assay of IRF5-high and -low 4T1 expressing cells. 5,000 cells were plated in 96-well plates and incubated for 12h. **(B)** Similar to **(A)** except cell proliferation was determined by alamarBlue™ assay. **(C, D)** Results from transwell migration **(C)** and invasion **(D)** assays are shown. Equal number of cells were plated in upper chambers and migration to serum-containing media allowed for 7h. Invasion assays were performed with Matrigel. Data are from 3 independent replicates. ****p≤ 0.0001. **(E)** Representative images from BLI of primary tumor growth and lung metastasis at the indicated time points post-fat pad injection of 0.1X10^6^ 4T1.2Luc3-IRF5-high (hi) or 4T1.2Luc3-IRF5-low (lo) cells. Images are representative of n=6 mice per group. **(F-H)** Primary tumor weights **(F)** and metastatic lung nodules **(G)** were determined at Day 30 post-injection. Representative lung images from **(G)** are shown in **(H)**. *p≤ 0.05, **p≤ 0.01. **(I)** Survival of mice implanted with 4T1-IRF5-high or 4T1-IRF5-low tumor cells, followed by primary tumor resection at day 10 post-implantation. *p≤ 0.05.

### 4T1 IRF5-high tumors are immunogenic due to distinct TIL recruitment

Studies have demonstrated the importance of the tumor immune microenvironment (TIME) in either promoting or inhibiting tumor growth and metastasis (Hanahan and Weinberg, 2011; Jin and Jin, 2020; Neophytou et al., 2021). Thus, identification of the mechanisms that regulate tumor immune cell infiltration are of immense interest in cancer biology. IRF5 is known to play a role in immune cell trafficking via the regulation of cytokines and chemokines (Ban et al., 2016; Brune *et al*., 2020; Feng *et al*., 2012; Pimenta *et al*., 2015; Takaoka et al., 2005; Yan *et al*., 2020). Although we previously reported in human BC that IRF5 directly regulates intra-tumoral cytokine and chemokine expression (Pimenta et al., 2015), the *in vivo* consequences of altered intra-tumoral IRF5 expression on immune cell recruitment have yet to be examined.

Thus, we investigated if intra-tumoral IRF5 expression alters the TIME. We found that IRF5-high 4T1 cells promoted increased tumor infiltration of CD8^+^ T cells, M1-like macrophages and CD11c^+^CD103^+^ tumor migratory dendritic cells (DCs) (**Fig. 8A-D**). In addition to increased numbers of infiltrating immune cells, we observed enhanced functional capacity through Perforin and Granzyme B staining of CD8^+^ T cells (**Fig. 8E**). Last, we detected significant reductions in the infiltration of M2-like macrophages and myeloid-derived suppressor cells (MDSCs) in the TIME of IRF5-high 4T1 tumors (**Fig. 8F-I**).

**Figure 8:**
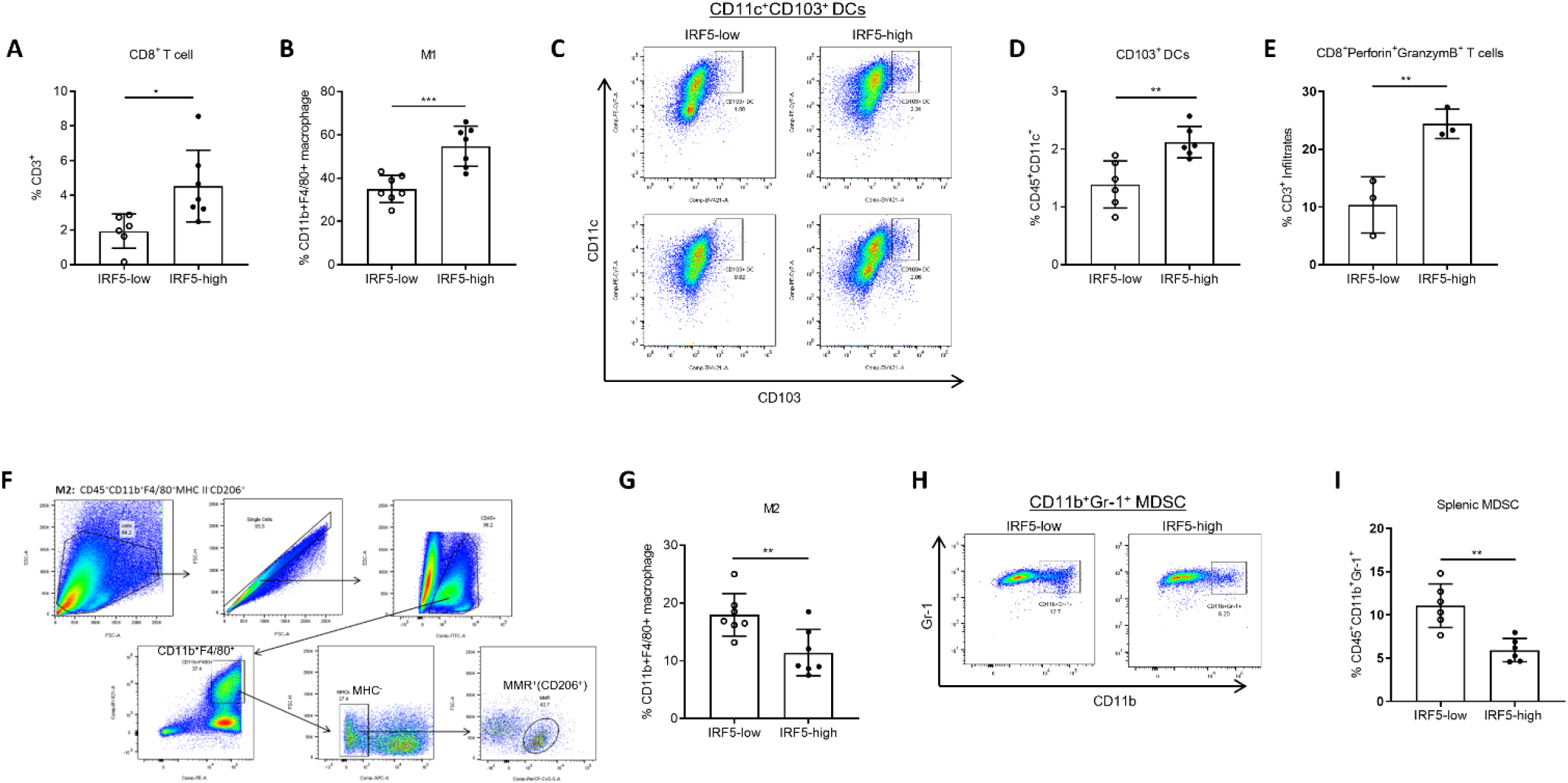
4T1-IRF5-high tumors are immunogenic due to distinct TIL recruitment. **(A,B)** Representative summary graphs of indicated TIL cell populations at Day 16 post-tumor cell implantation; n=6-7 mice/group. *p≤ 0.05, ***p ≤ 0.001. **(C,D)** Representative flow cytometry gating of infiltrating CD45^+^CD11b^-^ CD11c^+^CD103^+^ tumor dendritic cells **(C)** with quantification **(D)**; n=6 mice/group. **p ≤ 0.01. **(E)** Quantification of infiltrating CD8^+^Perforin^+^GranzymeB^+^ T cells; n=3 mice/group. **p ≤ 0.01. **(F,G)** Representative flow cytometry gating strategy for tumor infiltrating macrophages **(F)** with quantification of M2 macrophages **(G)**. **(H,I)** Representative gating strategy for splenic CD45^+^CD11b^+^CD11c^-^Gr-1^+^ myeloid-derived suppressor cells (MDSCs) **(H)** with quantification **(I)**.

### IRF5 expression is associated with increased TIL with elevated CD8A and CD14 expression

Elevated levels of TILs are associated with better clinical outcomes and response to neoadjuvant chemotherapy in TNBC (Asano et al., 2018; West et al., 2011). Previous work suggested that IRF5 expression in BC correlated with increased TIL recruitment (Garaud and Willard-Gallo, 2015; Pimenta and Barnes, 2014; Pimenta et al., 2015). To further explore this, we went back to the TCGA cohorts to examine correlations between *IRF5* BC expression and genes involved with TIL recruitment. As in our orthotopic mouse model, pathologic review of 161 H&E slides available from the TCGA-BIC cohort revealed increased TIL recruitment to IRF5-high (58%) versus IRF5-low IDCs (22%) (**Table I**, **Suppl. Fig. 8A, B**). A closer examination of TIL recruitment revealed that increased infiltration of multiple immune cell subsets including B and T cells, macrophages, and DCs, positively correlated with *IRF5* expression in TCGA-BRCA, Basal, Luminal, and Her2+ BC (**Suppl. Fig. 8B**). More specifically, we detected a positive correlation between *IRF5* expression and the abundance of *CD8A*, *CD14* and *CD11c* (*ITGAX*) expression in TIL, supporting increased infiltration of CD8^+^ T cells, CD14^+^ monocytes/macrophages and CD11c^+^ DCs, respectively (**Supp. Fig. 8C**). *IRF5* expression positively correlated with genes associated with increased CD8^+^ T cell infiltration such as *HLA-DMA* and *HLA-DMB,* T cell activation, and anti-tumor function, including *IFNG*, *CD40*, *LCK*, *SYK, GZMB* and *ICOS* (Callahan et al., 2008) (**Supp Fig. 8D, E**). We also found a significant positive correlation between *IRF5* expression and *CD86* expression in TILs that denotes B and T cell activation, dendritic cell activation and the M1 macrophage phenotype (**Supp. Fig. 8F**). Finally, we found that *IRF5* expression correlated with genes involved in immune cell migration, including *CCL4*, *CCL5*, *CXCR3* and *CXCL9* (**Supp. Fig. 8G**). These findings were both replicated and enhanced in basal-like TNBC (**Supp Fig. 9).** Taken together, these data reveal tumor intrinsic *IRF5* expression as a critical biomarker of the TIME in TNBC.

### Enrichment of ribosome biogenesis in mammary epithelial cells lacking Irf5

Our results demonstrate a protective role for IRF5 in mammary tumorigenesis both *in vitro* and *in vivo*. To further understand the mechanism(s) by which IRF5 regulates mammary tumor transformation, we performed RNA-seq on sorted luminal and myoepithelial cells from mammary glands of 6 weeks- and 9 months-old WT and KO mice. Epithelial cell purity in both age groups was confirmed with *Krt5* and *Krt8* expression (**Supp Fig. 10A**). IRF5 expression was also examined and confirmed to be higher in myoepithelial cells compared to luminal epithelial cells in both age groups (**Fig. 3B**). Gene set enrichment analysis (GSEA) results showed a relative enrichment in Epithelial to Mesenchymal Transition (EMT) and ribosome pathways in both pre-pubertal and aged KO mice (FDR q-value < 0.001) (**Fig. 9A, Supp Fig. 10B**). More detailed analysis of ribosomal transcripts revealed that 6 weeks-old KO mammary glands had higher levels of ribosomal subunit transcripts compared to their WT counterparts (FDR q-value < 0.001); similar albeit less dramatic findings were made in 9 months-old KO mice (**Fig. 9B**).

**Figure 9:**
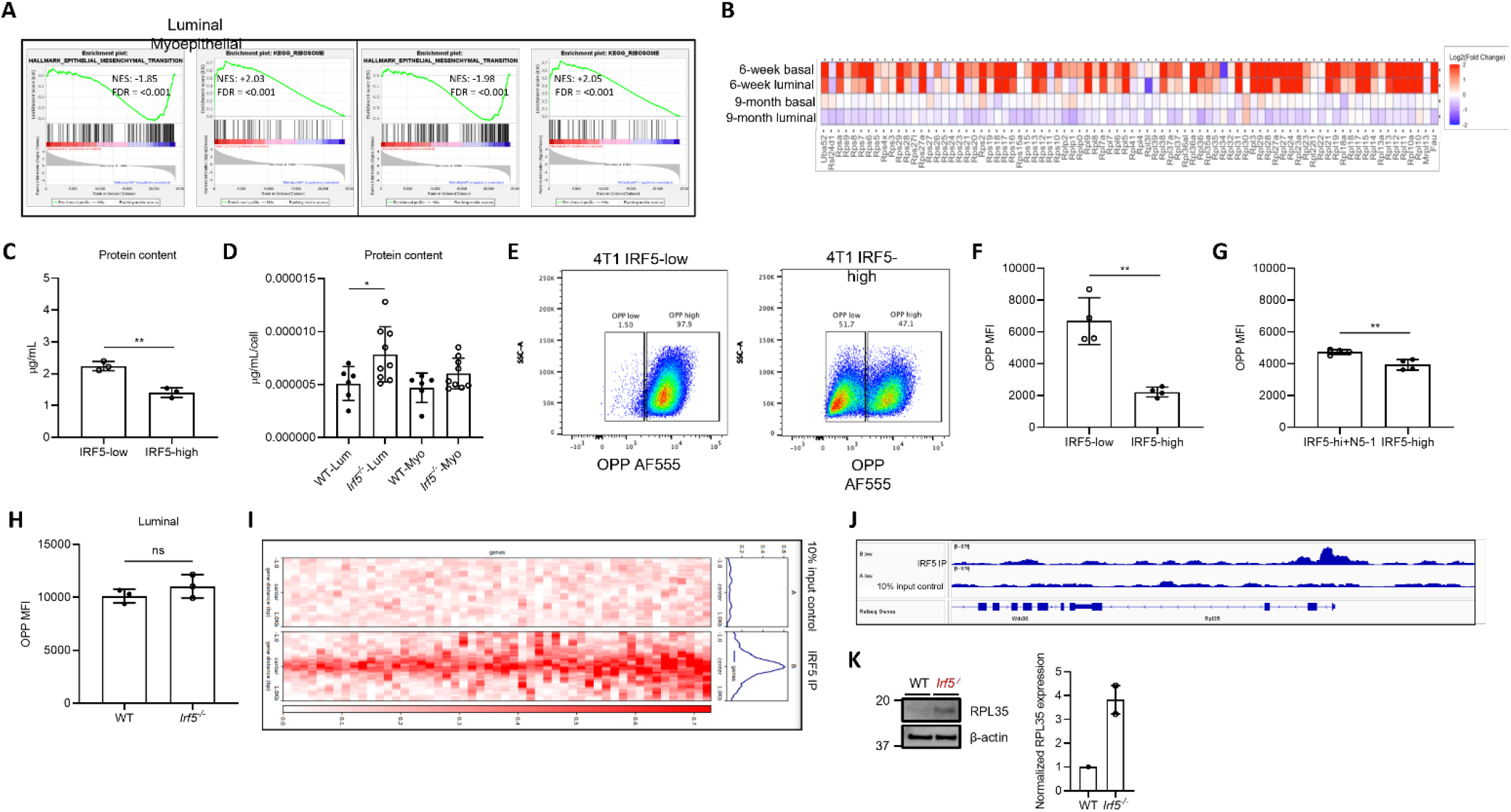
IRF5 regulates ribosome biogenesis in mammary epithelial cells. **(A)** GSEA of RNA-seq data from sorted luminal and myoepithelial cells of 9 months-old WT and *Irf5^-/^*^-^ mammary glands is shown. Representative terms were chosen with adjusted *p*<0.005. Red is *Irf5^-/-^* showing positive enrichment for KEGG_Ribosome (luminal, NES +2.03; myoepithelial, NES +2.05); blue is WT showing negative enrichment for HALLMARK_Epithelial to Mesenchymal Transition (luminal, NES -1.85; myoepthelial, NES -1.98). All FDR q-values <0.001. **(B)** Similar to **(A)** except a heatmap of the relative expression of genes involved in ribosomal biogenesis is shown as *Irf5^-/-^* compared to WT RNA transcript expression. **(C,D)** DC protein quantification assay in 4T-1 IRF5-high and -low cell lines **(C)** and primary sorted luminal and myoepithelial cells from WT and *Irf5^-/^*^-^ mice **(D)**. n=3 independent replicates are shown in **(C)**; n=6-9 mice/genotype are shown in **(D)**. *p≤ 0.05, **p ≤ 0.01. **(E- G)** Representative flow gating from OPP assay **(E)**. 4T-1 cell lines were plated on Day 0 at 100,000 cells/well in a 12-well plate and protein synthesis quantified on Day 1 **(F)**. **(G)** Similar to **(F)** except OPP assay was performed on 4T-1-IRF5 cells following treatment with the IRF5 inhibitor N5-1. n=4 independent replicates; **p ≤ 0.01. **(H)** OPP assay was used to measure rates of protein synthesis in primary sorted luminal mammary epithelial cells. (**I**) Gene enrichment analysis at all *Rpl* and *Rps* binding sites comparing to the control input sample from ChIP-seq. Mammary glands from n=6 WT mice were pooled for immunoprecipitation with anti-IRF5 antibodies. (**J**) Representation of IRF5 enrichment at the promoter region of *Rpl35* from ChIP-seq analysis. (**K**) Representative Western blot analysis and quantification of RPL35 protein levels in purified primary mammary epithelial cells from 6 weeks-old WT and *Irf5^-/-^* mice. Mammary glands from n=2-3 mice/genotype were pooled for each independent replicate.

Recent studies have shown that dysregulation of specific ribosomal subunits is highly correlated with cancer progression and metastasis (Ebright et al., 2020; Jaako et al., 2022; Liu et al., 2019; Yan et al., 2015; Zhao et al., 2021). A cursory review of the literature suggests that IRF5 has not yet been implicated in the regulation of ribosome biogenesis. Given the link between dysregulated ribosome biogenesis and tumorigenesis, we next performed DC assays on 4T1 cell lines and primary sorted cells from WT and KO mice to examine if IRF5 expression (high or low) impacted protein synthesis in mammary epithelial cells. We observed decreased protein content in IRF5-high compared to IRF5-low 4T1 cells (**Fig. 9C**). Quantitation of protein content in WT and KO sorted luminal and myoepithelial cells from 6 weeks-old mice demonstrated the similar finding of increased protein content in KO relative to WT luminal cells; similar trends were found in KO myoepithelial cells (**Fig. 9D**). We next examined if the observed increase in protein content in the IRF5-low 4T1 cells correlated with an increase in global protein synthesis using the O-propargyl-puromycin (OPP) assay. Indeed, IRF5-low 4T1 cells demonstrated a significant increase in protein synthesis compared to IRF5-high cells (**Fig. 9E, F**). To further confirm a direct role for IRF5 in protein synthesis, IRF5-high 4T1 cells were cultured in the presence of a validated peptide mimetic that inhibits IRF5 activation (Banga et al., 2020; Song et al., 2020). Select inhibition of IRF5 resulted in significantly increased rates of protein synthesis compared to mock-treated IRF5-high cells (**Fig. 9G**). Similar findings were made in KO luminal cells (**Fig. 9H**). Together, these results indicate a previously undescribed role for IRF5 in the regulation of protein translation in mammary epithelial cells.

### IRF5 is a direct regulator of ribosomal gene transcription in mammary epithelial cells

The protein levels of specific ribosomal subunits have been reported as either upregulated or downregulated during cancer development and metastasis (Ebright *et al*., 2020). To further understand the mechanism(s) by which IRF5 may alter ribosome biogenesis in mammary epithelial cells, we next determined the genomic binding pattern of IRF5 in an unbiased manner. We isolated total mammary epithelial cells from 6-8 weeks-old WT mice and performed chromatin immunoprecipitation with IRF5 antibodies followed by sequencing (ChIP-seq). In accordance with the RNA-seq findings, ChIP-seq analyses identified IRF5 as a global regulator of ribosomal transcription (**Fig. 9I**). In support of these findings, GO enrichment analysis of IRF5- associated genes included posttranscriptional regulation of gene expression, regulation of translation, and multiple gene families associated with the regulation of translation, mRNA and protein folding (**Suppl. Fig. 10B**). We next focused in on a ribosomal gene recently implicated in metastatic breast cancer. RPL35 expression in a mouse model of HR+ mammary tumorigenesis was shown to drive a significant increase in metastatic burden (Ebright *et al*., 2020). Results from ChIP-seq analysis revealed the direct binding of IRF5 at the predicted *Rpl35* promoter locus (**Fig. 9J**). We found RPL35 to be both transcriptionally (RNA-seq) and translationally upregulated in KO primary mammary epithelial cells (**Fig. 9K**). Thus, it is tempting to speculate that some of the observed IRF5-mediated alterations in the translational landscape of mammary tumor cells may be due to the transcriptional regulation of *Rpl35* by IRF5, which in turn contributes to the increased metastatic burden and poor clinical outcomes observed in IRF5-low or null breast cancers.

## Discussion

TNBC accounts for approximately 15% of all BC incidents and has the lowest rates of survival, occurring most frequently in women under 40 and often within 5 years of giving birth. Effective therapies guided by biomarkers that predict disease progression and response for patients with TNBC remains an unmet need. Here, analysis of BC data from TCGA revealed that IRF5 is a key biomarker of metastasis, TIL and survival. Follow up histologic and molecular studies of mammary glands from age-matched WT and KO littermate mice revealed new roles for IRF5 in the regulation of murine mammary gland development, involution, TIL regulation, tumor-associated ribosomal biogenesis, and protein translation. Further, many of our findings in mice are consistent with previously reported roles for IRF5 in the regulation of immune cell responses, apoptosis, and tumor suppression in human cells.

Hyperactivation of ribosomes and increased protein translation are considered critical for cancer initiation and progression (Gachet et al., 2018; Pelletier et al., 2018; Song et al., 2021). However, recent studies have revealed a more complex relationship between ribosome biogenesis and cancer. In 2008, studies of the proto-oncogene MYC revealed that MYC drives increased 7-methylguanosine cap-dependent mRNA translation over internal ribosome entry (IRE) translation, resulting in suppression of transcripts utilizing IRE sites including cyclin dependent kinase 11 (CDK11) (Petretti et al., 2006). In addition, increasing translation has been shown to upregulate genes implicated in ribosome biogenesis, which have in turn been linked to metastasis. As the largest and most transcriptionally active region in the human genome, the rDNA encoding ribosomes themselves are inherently unstable and a known loci for DNA damage (Salim and Gerton, 2019). Homologous recombination is the preferred double-stranded break repair mechanism of choice for the damage associated with repetitive sequences (Korsholm et al., 2019). However, in the context of Balb/c mice with reduced DNA-PK activity and impaired DSB repair machinery, we postulate that the increased transcription of rDNA driven by loss of *Irf5* creates the perfect storm for increased mutational burden (Okayasu et al., 2000). In the current study, we provide evidence for a previously unrecognized regulatory role for IRF5 in tumor-associated ribosome biogenesis and protein translation, where loss of *Irf5* expression results in a drastic upregulation of rRNA transcription and translation of the tumor-associated ribosomal subunit, RPL35. Thus, we propose that silencing of *IRF5* in human breast cancer may uniquely contribute to the “multi-hit” model of carcinogenesis through a two-fold mechanism; 1) by directly driving increased expression of tumor-associated ribosomal proteins as we demonstrate here and 2) by increasing the likelihood for somatic mutations through drastic global upregulation of unstable rDNA genomic loci transcription (Smirnov et al., 2021).

In addition to a role for IRF5 in the molecular regulation of ribosomal translation, we identified a particularly interesting role for IRF5 in mammary gland involution. Following involution induction, KO breeder mice had significant alterations in mammary gland remodeling and demonstrated increased rates of cancer with age. Interestingly, IRF5 was predicted to be activated during involution and mastitis infection (Sharp et al., 2016). In human BC, pregnancy is linked to a reduction in risk except in the case of the HER2+ subtype. It has been postulated that accumulation of pregnancy-identified mammary epithelial cells (PIMECs), a pregnancy derived epithelial subpopulation that persists post-pregnancy as multipotent stem cells, increases the risk of BC by acting as tumor-initiating cells upon loss of endogenous tumor suppressive functions (Eyermann et al., 2021). In our initial analysis of the luminal cell populations, with whom PIMECs have a theorized shared lineage, we saw an increase in luminal ductal cells in *Irf5^-/-^* mammary glands (Chang et al., 2014). Following subsequent *in vitro* involution induction, we observed altered remodeling of the post-lactational mammary glands. Together, these findings indicate a likely alteration in the luminal progenitor cell populations, which in turn may create an environment primed for malignant transformation. Mechanistically, this is likely due in part to the previously described pro-apoptotic role for IRF5, where loss of *Irf5* in the context of programmed cell death during mammary gland involution may drive increased PIMEC survival. In addition, the wound-healing environment of involuting mammary glands is a known risk factor for tumor development and metastasis, supporting our observations of prolonged mammary gland remodeling, delayed lymphocyte infiltration, and decreased M1 macrophages in tumor prone KO mice (Lyons et al., 2011).

Lymphocyte function and infiltration are key mechanistic drivers in BC generation and predictive of clinical progression. In early stage TNBC and HER2+ BC, retrospective studies have shown that patients with increased TILs have significantly improved clinical outcomes and increased pathological complete response (pCR). However, use of TILs as a predictive biomarker for disease progression and neoadjuvant response is still far from common in clinical practice. This is attributed to the challenge of uniform pathological scoring of TILs, which can frequently be complicated by tumor growth pattern heterogeneity and the difficulty of distinguishing tumor cells from stromal TILs. Thus, aside from the traditional grading, staging, gene recurrence scores, and hormone receptor analysis of early BC, there are currently few clear biomarkers predicting tumor prognosis (Savas et al., 2016). Here, we demonstrate that IRF5 expression in human BC can predict not only the degree of TIL infiltration, but also the subsequent clinical course. Additionally, we show that BC expression of IRF5, a transcription factor known in immune cells to regulate not just proinflammatory cytokine production, but also immune cell migration, has a strong correlation with tumor immune cell infiltration. Importantly, we establish that IRF5 expression in human BC positively correlates with increased TILs and improved clinical outcomes. Thus, characterization of IRF5 expression and function in both human and mouse mammary gland development and tumorigenesis provides highly relevant clinical insight into IRF5 as a novel biomarker of tumor progression and metastasis, as well as a potential predictor for tumor response to immune-based therapies.

## Acknowledgements

This work was supported in part by the National Cancer Institute (1R21CA1952561), DoD BCRP (W81XWH-19-1-0113), and the Manhasset Women’s Coalition Against Breast Cancer (BJB). LB was a recipient of the Allied World St. Baldrick’s Foundation career development award. We would like to acknowledge and thank Dr. Xiaohu ‘Emma’ Bi for the original generation of the colony of *Irf5^-/-^* and *Irf5^+/+^* Balb/c littermate mice.

## Author contributions

BJB and DL designed the study; ZB, DL, SS, QG, KK, ML (Lallo), ML (Lapan), VN and MR performed all *in vitro* and *in vivo* experiments characterizing *Irf5^-/-^* and *Irf5^+/+^* littermate mice; CS and DIL generated and characterized the 4T1 IRF5-high and low mouse models of mammary tumorigenesis and metastasis; IC, EP and RR performed analysis of TCGA data; DL, ZB, MM and COS performed RNA-seq and analysis; ZB, MR and MM performed ChIP-seq and analysis; ZB, DL, SS, KK, MR, MM, BJB, LB, COS and KP performed data analysis and generated the figures; ZB, DL, BJB, LB and KP prepared the manuscript.

## Methods

### Mice

*Irf5^+/+^* and *Irf5^-/-^* Balb/c mice were generated by crossing *Irf5^-/-^*C57Bl/6 mice to wild-type Balb/c mice purchased from The Jackson Laboratory. *Irf5^-/-^*Balb/c mice were then back-crossed for more than 12 generations to obtain a >98% pure background. Due to the nature of the study, only female age-matched *Irf5^+/+^* and *Irf5^-/-^*Balb/c littermate mice were used. For the 4T1 orthotopic model of TNBC, 8-12 weeks-old female wild-type Balb/c mice were used. Mice were kept in environment-controlled pathogen–free conditions with a 12-hour light/dark cycle and an ambient temperature of 23°C ± 2°C. This study was carried out in strict accordance with recommendations in *Guide for the Care and Use of Laboratory Animals* of the NIH (National Research Council, Guide for the Care and Use of Laboratory Animals (National Academies Press, Washington, DC, ed. 8, 2011). The protocol was approved by the Institutional Animal Care and Use Committee (IACUC) of the Feinstein Institutes for Medical Research and the USAMRDC Animal Care and Use Review Office (ACURO).

### Cell lines, plasmids, retroviral transduction and *in vitro* cellular assays

The murine 4T1 mammary carcinoma cell line and the Phoenix viral packaging cell line were purchased from American Type Culture Collection (ATCC; Manassas, VA, USA). The luciferase-expressing 4T1 (4T1.2-Luc3) cell line was provided by Dr. Cheryl Jorcyk (Boise State University). All cell lines were confirmed negative for pathogens by Radil^®^ testing (IDEXX BioAnalytics; Westbrook, ME, USA). Cells were cultured at 37°C in 5% CO_2_/95% air in Dulbecco’s modified Eagle’s medium (Sigma, St Louis, MO, USA) containing 10% heat-inactivated fetal bovine serum (Sigma). Full-length murine *Irf5* was cloned into the pBabepuromycin vector as described (Bi *et al*., 2011), using the BamHI/SalI sites. pBABE-Con (empty vector control) and pBABE-Irf5 plasmids were transfected into Phoenix packaging cells. Phoenix supernatants containing virus were collected at 24 and 48 h post-transfection and used to transduce 4T1 and 4T1.2-Luc3 cells. After 2 days, stable clones (4T1-Con, 4T1.2-Luc3-Con, 4T1-Irf5, and 4T1.2-Luc3-Irf5) were selected for using puromycin selection media and colonies pooled. Expression of Irf5 was confirmed by Western blot. Cell viability, proliferation, transwell migration and invasion assays were performed as previously described ((Bi *et al*., 2011; Pimenta and Barnes, 2015). Briefly, viability was determined by MTT assay (ThermoFisher Scientific, V13154) and proliferation with alamarBlue™ Cell Viability Reagent (ThermoFisher Scientific, DAL1025), as per manufacturer’s instructions.

### TCGA data collection and analysis

RNA-seq transcriptomic data from TCGA (BRCA: Cell 2015, Firehose legacy, Nature 2012, and PanCancer Atlas (n=2,990) was downloaded from the UCSC Xena portal and matched normal breast epithelial tissue (GSE86354) from the NCBI Gene Expression Omnibus (Ciriello et al., 2015; Edgar et al., 2002; Gao et al., 2013; Goldman et al., 2020; Hanahan, 2022; Hoadley et al., 2018; Koboldt et al., 2012; Pereira et al., 2016). Corresponding clinical data were retrieved from the same source. scRNA-seq data from normal breast epithelial (GSE113197; n= 867), primary tumor (GSE75688 & GSE118389; n=1,851) and CD45^+^ immune cells (GSE114725; n=21,253) was downloaded from (Aran et al., 2017). Immune cell infiltration was determined using TIMER 2.0 Immune Outcome module (Li et al., 2016; Li et al., 2017). The T cell infiltration gene expression signature includes the following 13 genes: CD8A, CCL2, CCL3, CCL4, CXCL9, CXCL10, ICOS, GZMB, IRF1, HLA-DMA, HLA-DMB, HLA-DOA, HLA-DOB (Spranger et al., 2015). Spearman correlation was used to quantify the association between IRF5 gene expression and immune cell expression. The association between IRF5 expression and survival was evaluated by Kaplan–Meier analysis (using the Kaplan-Meier plotter) (Győrffy, 2021; 2023; Lánczky and Győrffy, 2021). Samples were stratified in two groups according to their IRF5 gene expression (low and high) using the JetSet Best Probe Set (Li et al., 2011). The 25^th^ (low) and 75^th^ (high) percentiles were used as cutoff thresholds. The hazard ratios (HRs) were computed with 95% CIs and log-rank *p* values. The cBio Cancer Genomics Portal (cBio-Portal) was used to evaluate the somatic mutation and copy number variation (CNV) profile of IRF5 in TCGA-BRCA and METABRIC (Curtis et al., 2012).

### Protein quantification using BCA assay

The day prior to the assay, 500 x 10^3 4T1 cells expressing Irf5-luciferase or control vector were plated in triplicate in 24 well plates. The day of the assay, cells were trypsinized, washed once in PBS, and counted. 300,000 cells were then pelleted. Supernatant was discarded and pellets were lysed in 50µL of RIPA buffer supplemented with protease inhibitor cocktail. Lysates were diluted 1:10 in RIPA Lysis and Extraction buffer (Thermo Fischer) supplemented with protease inhibitor cocktail (Fischer Scientific) for use in BCA protein assay (BioRad). BCA protein assay was performed in triplicate for each diluted sample following manufacturer’s protocol. N = 3 independent experiments.

### Cell sorting and immunoblot analysis

Luminal and basal myoepithelial cells were prepared from 5 WT and 5 *Irf5^-/-^* 6-week-old murine mammary glands as previously described (dos Santos *et al*., 2013). After dissociation cells were stained with anti-mouse CD45-FITC, anti-mouse Ter119-FITC, anti-mouse CD31-FITC antibodies, anti-mouse CD24 eFluor 450 (Biolegend) and anti-mouse CD29-PE-Cy7 (Biolegend) for 30 minutes at 4C. Luminal and basal cell populations were then sorted using BDFACSAria and pooled for immediate immunoblot analysis.

Whole cell lysates were prepared by lysing cells in NP-40 lysis buffer (50 mM Tris-HCl (pH 7.4), 150 mM NaCl, 1% NP-40 and 5 mM EDTA) (Thermo Scientific, J60766-AP) supplemented with Halt Protease Inhibitor Cocktail (Thermo Scientific, 87786). Sample protein concentrations were quantified using the DC protein assay (Bio-Rad, 5000112). In general, 15-25 μg of protein per sample were separated by SDS-PAGE using the Bolt Bis-Tris system (Invitrogen). Proteins were transferred to 0.45 μm nitrocellulose membranes (MDI, SCNX8402XXXX101) using a wet transfer system. Transfer efficiency was assessed by incubating the membranes in 5 mL of Ponceau S (Sigma-Aldrich, P7170) for 5 min followed by distaining with Tris- buffered saline-0.05% Tween 20 (TBST). The membranes were blocked for 1 h at room temperature (RT) with 5% bovine serum albumin (BSA) in TBST and incubated overnight at 4°C with the primary antibody diluted in the blocking buffer. The membranes were washed three times for 5 min each with TBST and incubated with the secondary antibody diluted in the blocking buffer for 1 h at RT. The membranes were washed three times for 5 min each with TBST and incubated with 1 mL of chemiluminescent detection reagent (Cytiva, RPN2232) for 3 min before image acquisition. Horseradish peroxidase (HRP)-conjugated β-actin antibody (Cell Signaling, 12620, 1:5000) was used as the loading control for protein normalization. Densitometric analysis was performed using the Image Lab software (Bio-Rad).

### *In vivo* tumor initiation, metastasis and survival

To initiate primary tumor growth and lung metastasis, freshly prepared 1 x 10^5^ 4T-1-con, 4T-1-Luc3-Con, 4T-1-Irf5, or 4T-1-Luc3-Irf5 cells in mid-log phase growth were orthotopically injected into the 4^th^ right mammary fat pad of 8-12 weeks-old female Balb/c mice (Jackson Laboratory, Bar Harbor, ME). The volume of primary tumors was quantified weekly using digital calipers by measuring in three dimensions. 4T-1-Luc3 tumors and metastases were monitored by bioluminescence imaging on an IVIS Lumina III System (Perkin Elmer) twice a week after tumor implantation. When primary tumors reached 15 mm in the largest diameter, mice were sacrificed, tumors weighed, and organs harvested for fixation and histopathological analysis or single-cell preparation and flow cytometry analysis of immune cell composition. Lung metastases were quantified by counting the number of visible nodules on the surface of the lung. Counting was performed in a blinded manner by 3 independent readers. In a separate cohort of mice, survival studies were performed after primary tumor resection at day 10 post-tumor cell injection. Animals were sacrificed when moribund and metastasis determined by visual inspection.

### Flow cytometry analysis

Flow cytometry was carried out on single-cell suspensions of mammary glands. A standard protocol was used to prepare single-cell suspensions for flow cytometry analysis of luminal and myoepithelial cell populations as previously described(dos Santos *et al*., 2013). For intracellular staining, cells were fixed and permeabilized, after surface staining, with the FoxP3 Fixation/Permeabilization kit (eBioscience) according to the manufacturer’s instructions, followed by appropriate intracellular staining. Flow cytometric acquisition was completed using LSR-Fortessa (BD Biosciences), and analysis was performed using FlowJo (Tree Star).

### RNA *in situ* hybridization

Sections from formalin-fixed paraffin-embedded (FFPE) mammary glands of WT mice were stained following the manufacturers protocol (RNAscope^®^ 2.5 HD Duplex Reagent Kit, Advanced Cell Diagnostics (ACD Cat No: 322436)). RNAscope Target Probe for murine *Irf5* was used. ACD’s universal negative control targeting the *dapB* gene (GenBank accession #EF191515) was used to assess specificity and the positive control probe gene Cyclophilin B (*PPIB*) was used to assess tissue and RNA quality.

### 3D *in vitro* culture of mammary epithelial organoids

Primary mammary organoids were derived from inguinal mammary glands of 6 weeks-old and 9 months-old female WT and KO nulliparous mice, as previously described (Ciccone et al., 2020; Ewald, 2013; Lewis et al., 2022). Lymph nodes were removed. Mammary glands were minced with slides and incubated in DMEM/F12 supplemented with 5% FBS and gentle collagenase/hyaluronidase (StemCell Catalog #07919). After centrifugation (2000 r.p.m., 5 min), the pellet was resuspended in DMEM/F12 and treated with DNase 1 (20 U/mL). Epithelial pieces were separated from the single cells through a series of differential centrifugation. The final pellet containing organoids was re-suspended in Matrigel and distributed in 96-well culture plates containing an underlay Matrigel layer (Growth Factor Reduced Matrigel, BD Biosciences, San Jose, CA, USA). Cultures were maintained at 37 °C in serum-free basal medium (DMEF/F12, 1% ITS (insulin/transferrin/selenium), 1% penicillin/streptomycin), BOM medium was changed every three days. Lactation media was changed every two days. Colonies were imaged and counted using the EVOS M7000 imaging system.

### Whole-mount analysis of mammary glands

Carmine-alum staining was performed as described (<u>19).</u> Briefly, dissected mammary glands were spread onto positively charged glass slides, fixed in 10% Neutral Buffered Formalin overnight, hydrated, stained overnight in 0.2% carmine, dehydrated in graded solutions of ethanol, cleared in toluene overnight and mounted. The slides were scanned, and digital pictures taken on the EVOS M7000. Filled fat pat area was measured as described (Pruitt et al., 2018; Zuo et al., 2014). The invasion distance or lymph node area was scored by measuring from the lymph node to the farthest epithelial TEB (Zuo *et al*., 2014). Number of TEBs were counted. These methods are widely used in mammary development biology (Xiong et al., 2020).

### Histology and immunohistochemistry

For histologically based assays, the inguinal glands were dissected, fixed in 10% neutral buffered formalin (Anatech Ltd.) for 24 hours, then transferred to 70% EtOH prior to processing. All samples were then sent to Histowiz for paraffin embedding,4 μm sectioning, H&E staining, and blinded analysis. Antigen retrieval prior to IHC and subsequent staining was performed as previously described (Abcam).

### Experimental mice and reproductive staging

All experiments were conducted using pubertal (5 - 6-week-old), breeder (20 - 34 weeks) or old mice (9 - 18 months old). To initiate involution, pups were weaned between 9 and 13 days of age. Mice were euthanized on designated days of involution and left and right inguinal and thoracic mammary glands harvested.

### RNA-Seq and ChIP-Seq

Luminal epithelial and myoepithelial cells were sorted from three mice per age and genotype. RNA extraction was performed with the RNAeasy mini kit (Qiagen). Double stranded cDNA synthesis and Illumina libraries were prepared utilizing the Ovation RNA-seq system (V2) (Nugen Technologies, #7102-32). RNA-seq libraries were prepared utilizing the Ovation ultralow DR multiplex system (Nugen Technologies, #0331-32). Each library was pooled and barcoded with Illumina True-seq adaptors to allow sample multiplexing, followed by sequencing (n=4) on an Illumina NextSeq500, high output, 76bp single-end run. Data quality was assessed using the FastQC software (Andrews, 2010). Adapter removal and quality trimming was performed using the Trimmomatic software tool with recommended settings for single end reads (Bolger et al., 2014). Trimmed reads were aligned to the mm9 reference genome (GenBank accession GCA_000001635.1) using STAR version 2.6.0a, a splice aware short read alignment software (Dobin et al., 2012). Read count quantification was performed using the FeatureCounts tool in the RSubread package (Liao et al., 2019). Differential expression analysis was performed using the DESeq package (Anders and Huber, 2010) in R (Team, 2022). Gene set enrichment analysis was performed using the GSEA tool version 2.1.0 (Mootha et al., 2003; Subramanian et al., 2005), and heatmaps were generated using the Heatmap2 function of the gplots package (Warnes et al., 2022) in R.

Chromatin immunoprecipitation was performed using the Pierce Magnetic ChIP Kit (Thermo Scientific, 26157). Briefly, 15 x 10^6^ primary mammary epithelial cells were sorted from 6 WT Balb/c mice (6-8 weeks-old). The cells were crosslinked with 1% formaldehyde for 10 min at RT and quenched with a glycine solution. The cells were washed three times with 4 mL of ice-cold phosphate-buffered saline (PBS) supplemented with EDTA-free Halt Protease and Phosphatase Inhibitor Cocktail (Thermo Scientific, 78441) and lysed with Membrane Extraction Buffer for 10 min on ice. The nuclei were harvested and incubated with 8U of micrococcal nuclease (MNase) in MNase Digestion Buffer Working Solution for 15 min at 37°C. The digestion was quenched via the addition of MNase Stop Solution, and the nuclei were harvested and resuspended in IP Dilution Buffer. The sample was sonicated for three 20-second pulses at 6 watts power (Sonicator 3000) for nuclear disruption. The digested chromatin was recovered and 10% of the sample was set aside as the 10% input sample. The remaining chromatin was incubated with 6 ug of anti- IRF5 antibody (Bethyl Laboratories, A303-386A) overnight at 4°C with mixing for IRF5 immunoprecipitation. The IP reaction was incubated with ChIP Grade Protein A/G Magnetic Beads for 2 h at 4°C with mixing and then the beads were washed with IP Wash Buffers 1 and 2. The beads were incubated in IP Elution Buffer for 40 min at 65°C with vortexing every 10 min. The 10% input and IRF5 IP samples were incubated with Proteinase K for 1.5 h at 65°C. The chromatin was recovered via Clean-Up Column purification and DNA integrity was assessed using a NanoDrop spectrophotometer.

Initial ChIP-seq library QC and alignment were performed by Novogene. Reads were aligned to the mm10 mouse reference genome. Following peak calling, Motif enrichment was performed using the Meme software suite, specifically the AME tool for Motif finding (McLeay and Bailey, 2010). Gene ontology analysis was performed using the Gene Regulatory Enrichment of Annotations Tool (GREAT) analysis web interface (McLean et al., 2010; Tanigawa et al., 2022). We specifically analyzed results from the Biological Processes and Cellular Function databases. Bigwig files for visualization were generated using the bamCoverage function in the NGS Deeptools software suite (Ramírez et al., 2016), and coverage heatmaps were generated using the multiBigwigSummary and plotHeatmap functions. Individual peaks were visualized in the Integrative Genomics Viewer genome browser (Robinson et al., 2011).

### *In vitro* protein synthesis quantification

OPP staining was performed using Click-iT plus OPP Alexa-555 protein synthesis kit per manufacturer’s protocol (ThermoFischer, Cat#: C10456). Briefly, 2 x 10^6^ cells were incubated at 25°C for 30 minutes with 1:400 dilution of the Click-iT OPP reagent. Cells were then washed three times with PBS and then fixed in 4% formaldehyde solution. Following fixation, cells were permeabilized in PBS supplemented with 2.0% saponin and 3.0 % FBS (v/v) for 15 minutes. For the Click-iT reaction, cells were incubated in the dark at room temperature for thirty minutes in the Click-iT reaction cocktail. Samples were then washed twice with PBS supplemented with 2% FBS and immediately analyzed. For samples using IRF5 peptide inhibitor - IRF5 N5-1 peptide inhibitor was added on Day 1 to a final concentration of 1 uM. Cells were harvested 24 hours later at 80% confluence. OPP assay was performed as per manufacturer recommendation to measure rate of protein synthesis.

### qRT-PCR

RNA was isolated from sorted mammary epithelial cells using the Qiagen RNeasy Mini Kit (Cat#: 74106). cDNA synthesis was performed using the GoScript reverse transcription system (Cat #: A5001). qPCR was performed in triplicate for each sample using the PowerUp SYBR green real-time PCR master mix with 5 – 10 ng input cDNA (cat#: A25776). Threshold values (*C*_T_) were averaged across sample replicates, followed by normalization *via* the ΔΔ*C*_T_ method to β-actin.

### Statistical analysis

Quantitative data are presented as mean ± SEM. Non-parametric data were analyzed by two-tailed Mann– Whitney *U* tests. Parametric data were analyzed using ANOVA with *post hoc* comparison (Tukey method). Adjusted *p*<0.05 was considered statistically significant.

### Immunoblot antibodies (Vendor, Catalog number, primary dilution, secondary dilution)

RPL35 (Proteintech, 14826-1-AP, 1:1000, 1:10,000)

IRF5 (Abcam, ab181553, 1:1000, 1:10,000)

Loading control: β-actin HRP conjugate (Cell Signaling, 12620, 1:5000)

Secondary: IgG HRP conjugate (Cell Signaling, 7074)

### Flow cytometry antibodies (Vendor, Catalog number, stock concentration, dilution)

FITC anti-mouse Ter119 (Biolegend, #116206, 0.5 mg/ml, 1:200)

FITC anti-mouse CD31 (Biolegend, #102406, 0.5 mg/ml, 1:200)

FITC rat anti-mouse CD45 (Biolegend, #103108, 0.5 mg/ml, 1:800)

CD45 anti-mouse PeCy7 (Biolegend, #103114, 0.2 mg/mL, 1:200)

PE-Cy7 anti-mouse/rat CD29 (Biolegend, #102222, 0.2 mg/ml, 1:100)

APC anti-mouse CD133 (Biolegend, #141208, 0.2 mg/mL, 1:100)

FITC anti-mouse/rat CD61 (Biolegend, #104306, 0.5 mg/mL, 1:100)

Pacific Blue™ anti-mouse/rat CD29 (Biolegend, #102224, 0.5 mg/ml, 1:250)

APC rat anti-mouse CD24 (Biolegend, #101814, 0.2 mg/ml, 1:250)

PE/Cy5 anti-mouse CD24 (Biolegend, #101812, 0.2 mg/ml, 1:250)

PE anti-mouse CD24 (Biolegend, #101807, 0.2 mg/ml, 1:250)

eF450 anti-mouse CD24 (ThermoFischer, #48024282, 0.2 mg/ml, 1:200)

Alexa Fluor® 700 anti-mouse CD24 (Biolegend, #101836, 0.5 mg/ml, 1:100)

Alexa Fluor® 647 anti-human/mouse CD49f (Biolegend, #313609, 1:100)

PE-Cy7 anti-human/mouse CD49f (Biolegend, #313622, 1:250)

PE anti-mouse Cd1d (Biolegend, #140805, 0.2 mg/mL, 1:200)

Live/Dead Fixable Yellow Dye (ThermoFischer, #L34959, 1:800)

Anti-CD16/CD32 Rat anti-mouse Fc Block (BD Pharmingen, #553142, 0.5 mg/mL, 1:400)

### Tissue staining antibodies (Vendor, Catalog number, stock concentration, dilution)

CK5 (Biolegend, #905501, 1.0 mg/mL, 1:2000)

CK8 (TTRO, Abcam, ab192467, 1:250)

α-Smooth Muscle Actin (Cell Signaling, #19245T, 1:500)

IRF5 (Abcam, ab181553, 1.189 mg./mL, 1:1000)

Ki67 (Abcam, ab15580, 1.0 mg/mL, 1:200)

### Primers

Keratin 8 (F-TGCAGAACATGAGCATTC, R-CAGAGGATTAGGGCTGAT)

Keratin 5 (F-AGTGCAGGCTGAGTGGGGAA, R-ACCAGCAAAGCCGCTGCCAAG)

