## Supplemental Data for "Loss of *IRF5* increases ribosome biogenesis leading to alterations in mammary gland architecture and metastasis"

### Supplemental Figure Legends

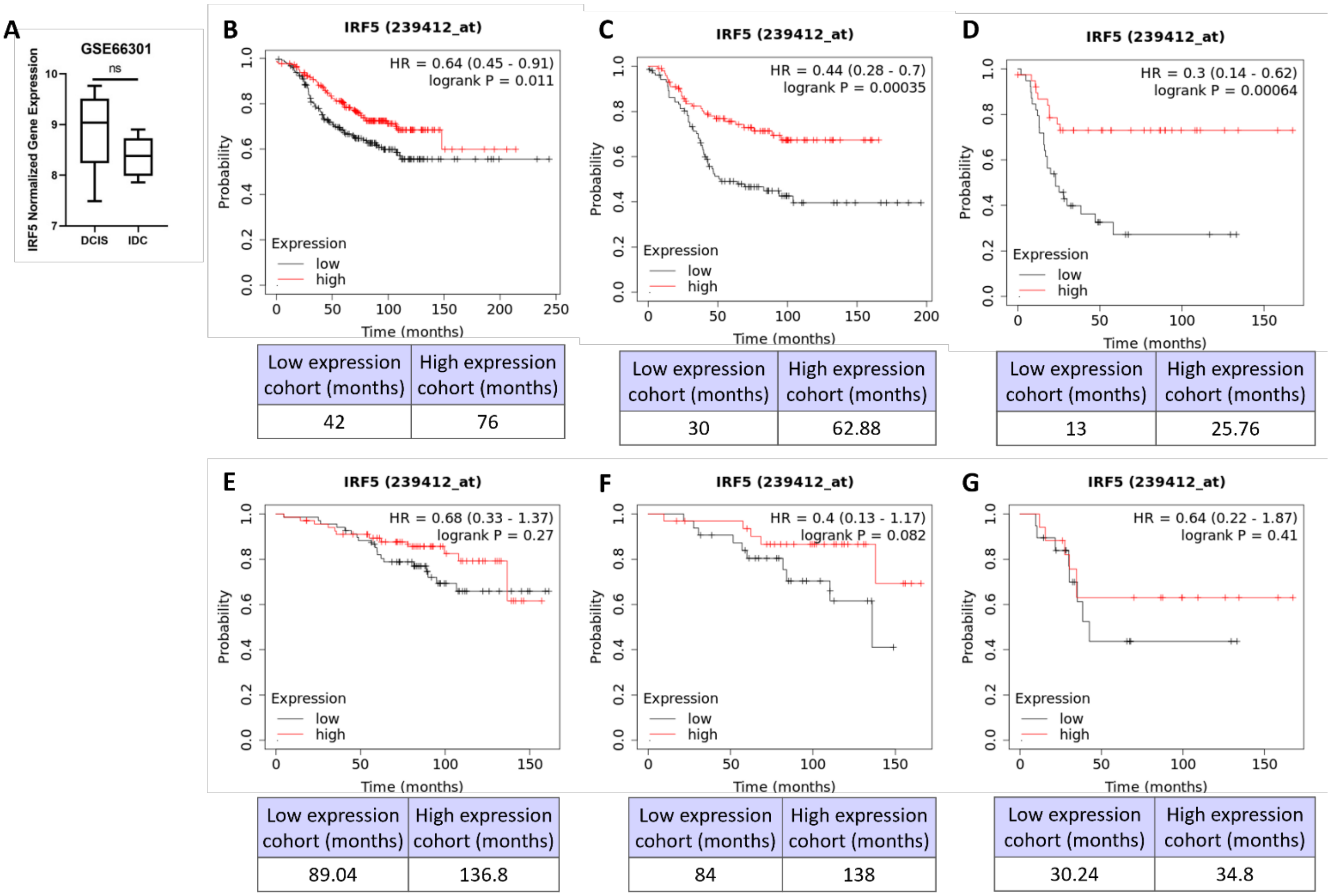


**Supp. Fig. 1. Loss of IRF5 expression is a marker of IDC and associates with poor RFS and OS in multiple BC subtypes.**

**(A)** Boxplot representation of *IRF5* expression within tandem DCIS/IDC patient samples ($n$ = 6) from GSE66301; normalized by the contributors (add 1, log2 transform, quantile normalization). Tandem DCIS/IDC were defined radiologically as DCIS lesions that have concurrent IDC within the same breast. Biopsy cores were subjected to RNA sequencing. ns = not significant.

**(B – D)** Data of RFS are from n=841 patients with luminal A BC (**B**), n=407 with luminal B BC (**C**), and n=156 with Her2+ BC (**D**) from the TCGA-BRCA cohort. Black line is lower quartile of IRF5 expression, red line is upper quartile. Graphs from Kaplan-Meier Database; JetSet best probe set used (IRF5 239412_at).

**(E – G)** Same as **(B – D)** except data are of OS from n=271 patients with luminal A (**E**), n=129 with luminal B (**F**), and n=73 with Her2+ (**G**).

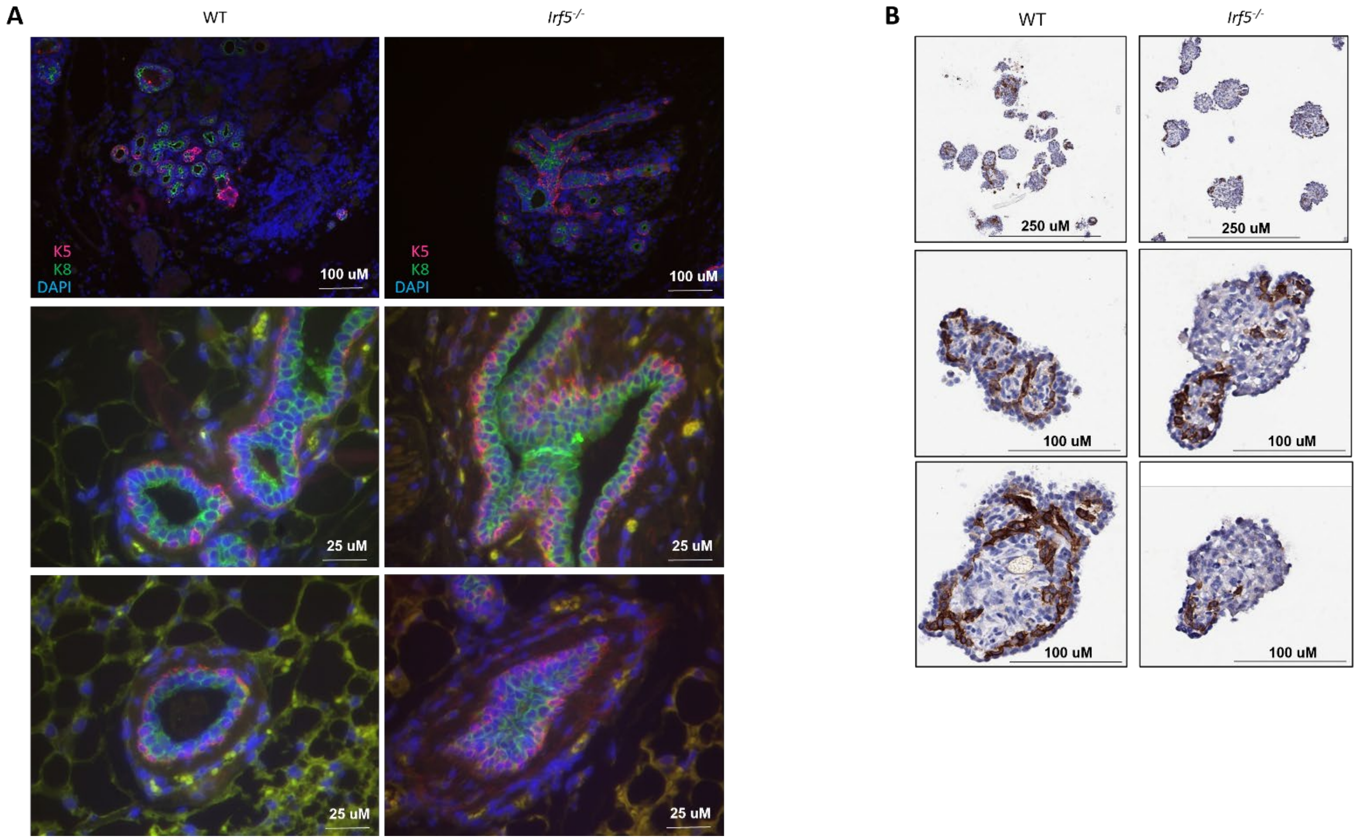


**Supp. Fig. 2.** Loss of IRF5 expression increases luminal cell proliferation

**(A)** Same as in **Fig. 2D**; representative IF staining of 12-month WT and *Irf5*$^{-/}$ mammary glands for K5 (basal; red), K8 (luminal; green) and DAPI (blue) demonstrating expansion of the luminal epithelial cell compartment.

**(B)** Representative SMA-IHC (brown) with hematoxylin counterstaining of mammary gland organoids from 9 months-old WT and *Irf5*$^{-/-}$ mice revealing a relative and select expansion of luminal epithelial cells in *Irf5*$^{-/-}$ mammary organoids; n=3/genotype performed in triplicate. Smooth muscle actin (SMA) is a sensitive marker of myoepithelial differentiation.

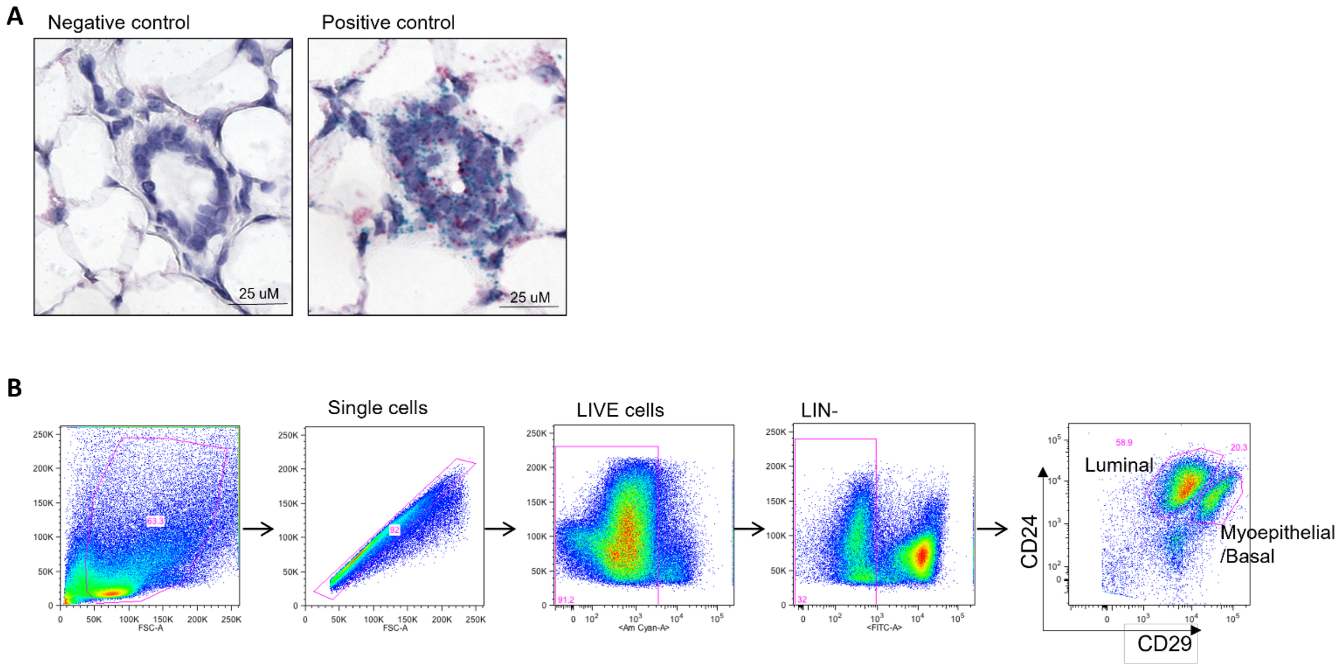


**Supp. Fig. 3.** Controls for murine *Irf5* ISH

**(A)** From **Fig. 3A**, ACD's universal negative control targeting the *dapB* gene (GenBank accession #EF191515) is shown to confirm *Irf5* target probe specificity in WT mammary glands and the positive control probe gene Cyclophilin B (*PPIB*) was used to assess tissue and RNA quality. Images were taken at 40X magnification.

**(B)** Representative flow cytometry gating of luminal and myoepithelial cell populations from 6 weeks-old WT mammary glands. Lin- depletion (Ter119$^-$CD45$^-$CD31$^-$).

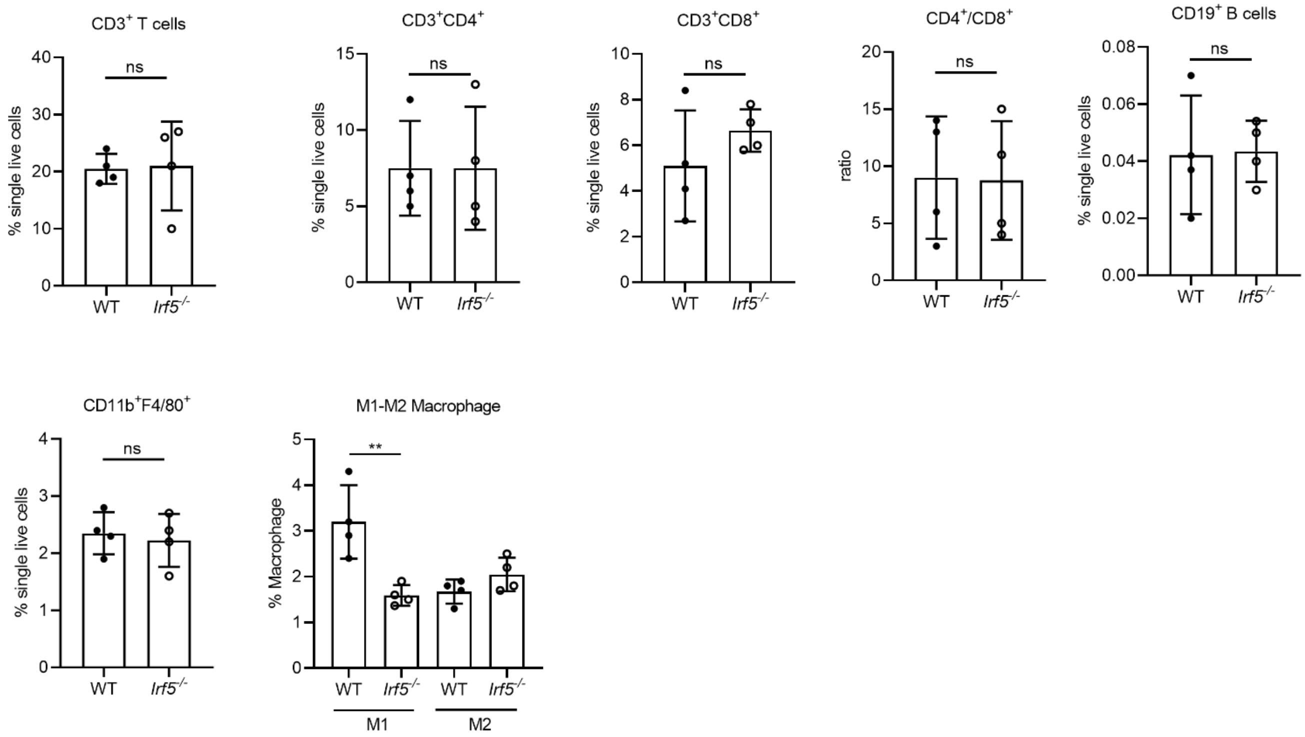


**Supp. Fig. 4.** Immune cell analysis in mammary glands of 9 months-old WT and *Irf5*$^{-/-}$ mice by flow cytometry.

Gated from $CD45^+$ cells. Percentage of infiltrated T cells, ratio of $CD4^+/CD8^+$ T cells, B cells, macrophages, and M1/2 macrophages is shown. ***p*< 0.001; ns = not significant. n = 4/genotype.

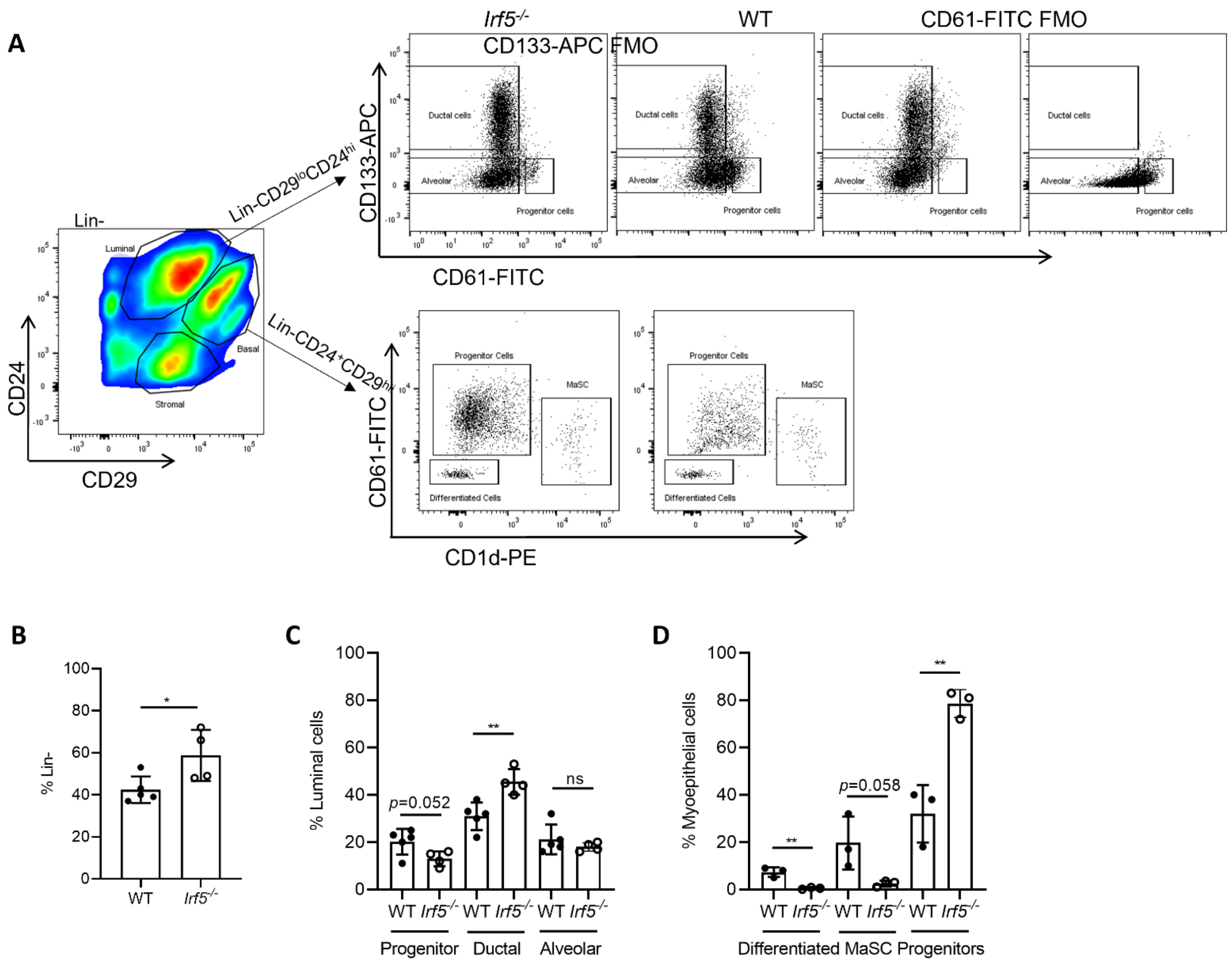


**Supp. Fig. 5.** Flow cytometry gating strategy for myoepithelial and luminal mammary cell subsets

**(A)** Representative gating strategy after Lin depletion (Ter119$^-$CD45$^-$CD31$^-$); shown in **Suppl. Fig. 3B)**. using CD24 and CD29 for detection of myoepithelial (basal), luminal and stromal cells within the mammary gland. Representative gating strategy for luminal ductal (Lin$^-$CD29$^{lo}$CD24$^{hi}$CD61$^-$CD133$^+$), alveolar (Lin$^-$CD29$^{lo}$CD24$^{hi}$CD61$^-$CD133$^-$) and progenitor (Lin$^-$CD29$^{lo}$CD24$^{hi}$CD61$^+$CD133$^-$) cells from age-matched WT and *Irf5*$^{-/-}$ mice.

**(B)** Quantification of luminal cells (Lin$^-$CD29$^{lo}$CD24$^{hi}$) from 6 weeks-old WT and *Irf5*$^{-/-}$mammary glands. *$p<$ 0.05; n=4-5/genotype.

**(C, D)** Quantification of luminal **(C)** and myoepithelial **(D)** cell populations from 6 weeks-old WT and *Irf5*$^{-/-}$ mammary glands. Myoepithelial cells were defined as Lin$^-$CD29$^{hi}$CD24$^+$. Myoepithelial differentiated cells (Lin$^-$CD29$^{hi}$CD24$^+$CD61$^-$), myoepithelial progenitor cells (Lin$^-$CD29$^{hi}$CD24$^+$CD61$^+$), and mammary gland stem cells (MaSCs, Lin$^-$CD29$^{hi}$CD24$^+$CD49f$^{hi}$CD61$^-$). **$p<$ 0.01, ns=not significant; n=3-5/genotype.

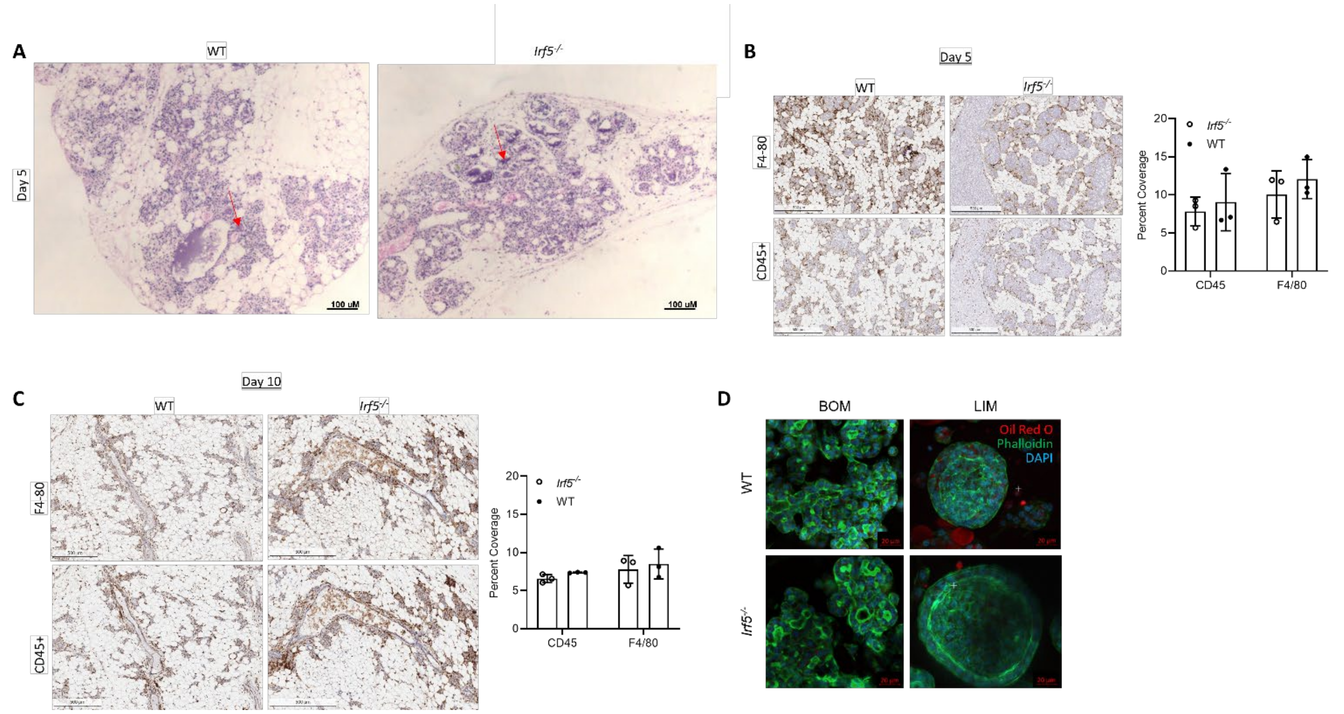


**Supp. Fig. 6.** Delayed or defective clearance of luminal milk producing cells in *Irf5*$^{-/-}$ mammary glands and mammary organoids

**(A)** Representative H&E staining of WT and *Irf5*$^{-/-}$ mammary glands at Day 10 post-involution induction showing retention of milk production (red arrows) in *Irf5*$^{-/-}$ mammary glands. Scale bar represents 25 uM.; n=3 mice per genotype.

**(B)** Representative CD45 and F4/80 IHC staining (left) of mammary glands with lymph node at Day 5 post-involution. Quantification is shown to the right. Scale bar represents 500 uM; n=3 mice per genotype.

**(C)** Same as **(B)** except mammary gland with lymph node is shown at Day 10 post-involution. Scale bar represents 500 uM n = 3 mice per genotype.

**(D)** Representative confocal images of WT and *Irf5*$^{-/-}$ organoids prepared from the mammary glands of 10 weeks-mice differentiated in either Basic Organoid Media (BOM) or Lactation Inducing Media (LIM). Images were taken 5 days following involution induction and then stained and analyzed by confocal microscopy. Red = Oil red O, Green = Phalloidin, Blue = DAPI. From n = 3 independent organoid cultures. Scale bar represents 20 uM.

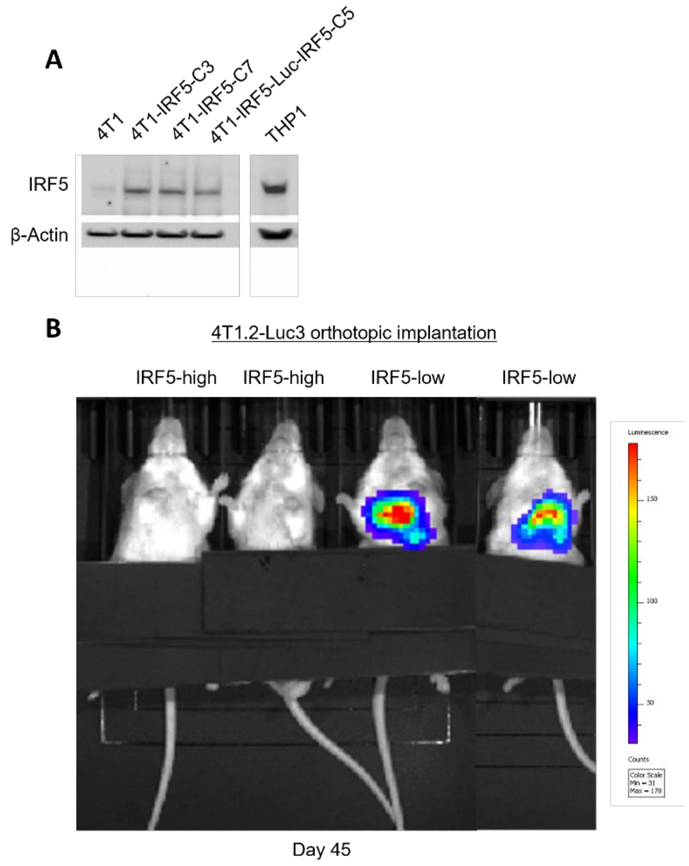


**Supp. Fig. 7.** IRF5 expression in 4T1 and 4T1.2-Luc3 cell lines

**(A)** Representative immunoblot of endogenous IRF5 expression in parental 4T1 cells and ectopic re-expression in 4T1 and 4T1.2-Luc3 cell lines. Positive control THP1 monocytes are shown.

**(B)** Live bioluminescence imaging (BLI) of lung metastases following orthotopic implantation of IRF5-high or -low 4T1.2-Luc3 cells into mammary fat pads of female Balb/c mice. Image taken at Day 45 post-implantation.

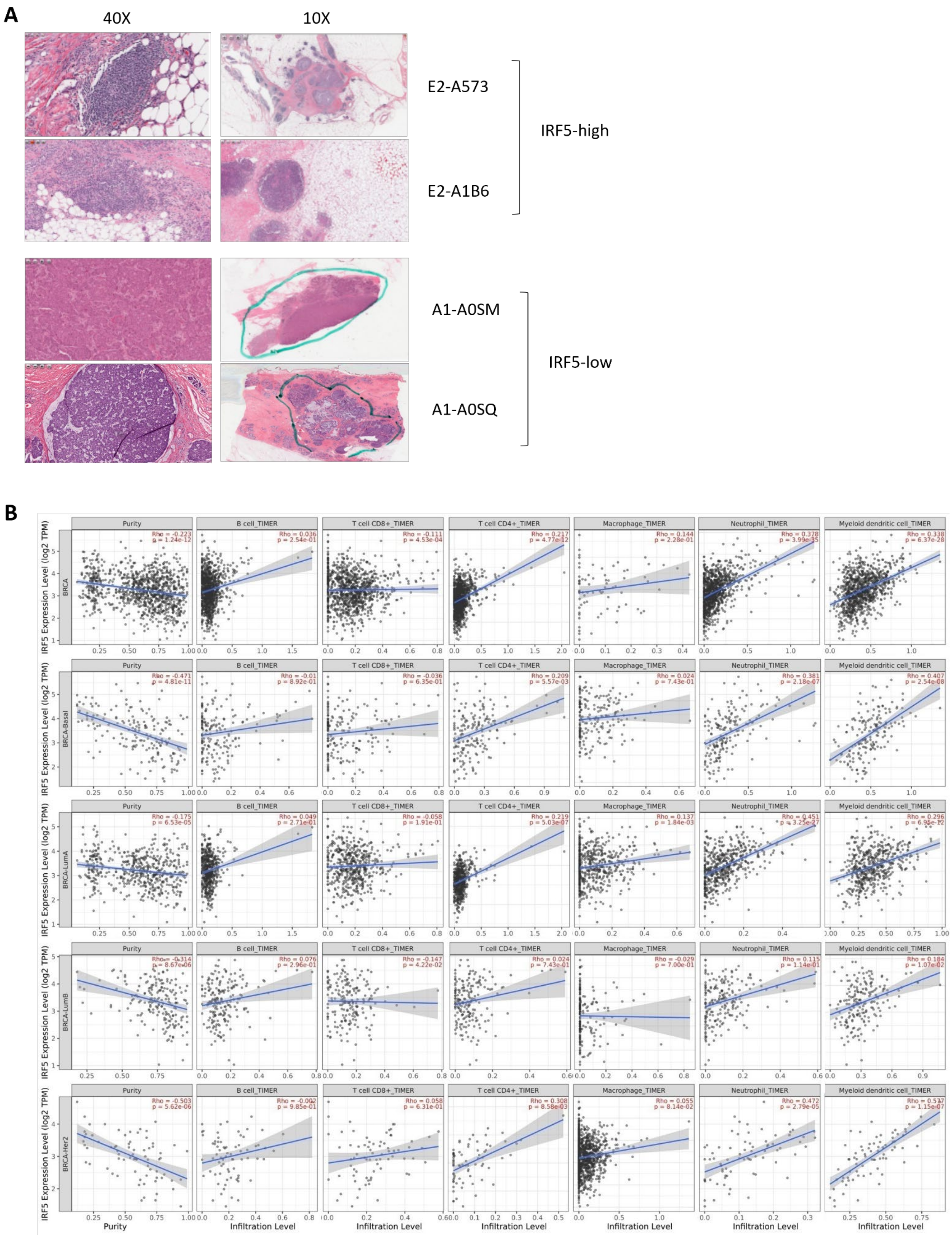
A
40X
10X
E2-A573
E2-A1B6
IRF5-high
A1-A0SM
A1-A0SQ
IRF5-low
B
Purity
B cell_TIMER
T cell CD8+_TIMER
T cell CD4+_TIMER
Macrophage_TIMER
Neutrophil_TIMER
Myeloid dendritic cell_TIMER
BRCA
BRCA-Basal
BRCA-LumA
BRCA-LumB
BRCA-Her2
IRF5 Expression Level (log2 TPM)
Infiltration Level

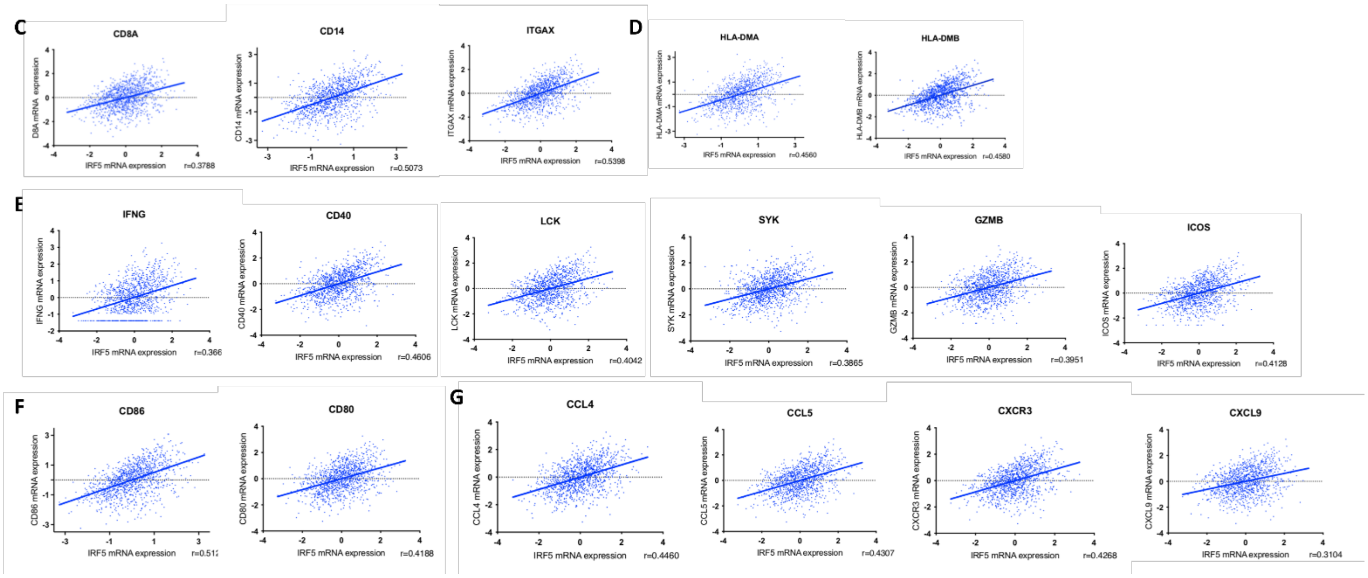


**Supp. Fig. 8.** *IRF5* expression correlates with increased TIL recruitment within the TCGA-BRCA and -BIC cohorts

**(A)** Representative H&E slides of IRF5-high (E2-A573 and E2-A1B6) and IRF5-low (A1-A0SM and A1-A0SQ) tumor tissue samples from the TCGA-BIC cohort (n=1105 patient samples. 6% (58/971) of the samples were IRF5-high and 94% were IRF5-low (Table I). E2-A573 and E2-A1B6 are from IRF5-high tumor specimens. E2-A573 shows several organized TIL structures. Right panel shows whole tissue sample, left shows single organized TIL. Pathology report states – invasive ducal carcinoma with lymphoplasmacytic infiltrate and necrosis. E2-A1B6 shows similar organized TIL structures. Pathology report states – ductal carcinoma in situ, solid type, nuclear grade 3 with necrosis, microcalcifications, and associated lymphoid infiltrate. A1-A0SM and A1-A0SQ are from IRF5-low tumor specimens. Pathology reports read – virtually no inflammatory response and no lymphocytic infiltrate for these samples.

**(B)** Correlation graphs from TIMER revealing increased TIL recruitment to IRF5-high versus IRF5-low tumors (y-axis), including all breast cancers (BRCA), Basal, Luminal A, Luminal B, and Her2+. Infiltration of B cells, $CD8^{+}$ T cells, $CD4^{+}$ T cells, Macrophages, Neutrophils, and Myeloid dendritic cells are shown. (x-axis). Data are from the TCGA PanCancer Atlas, female (n=1072).

**(C-G)** *IRF5* expression correlation analyses were performed using cBioPortal (TCGA PanCancer Atlas, all breast cancers, n=1072) with the indicated TIL markers and genes involved in tumor-immune cell activation.

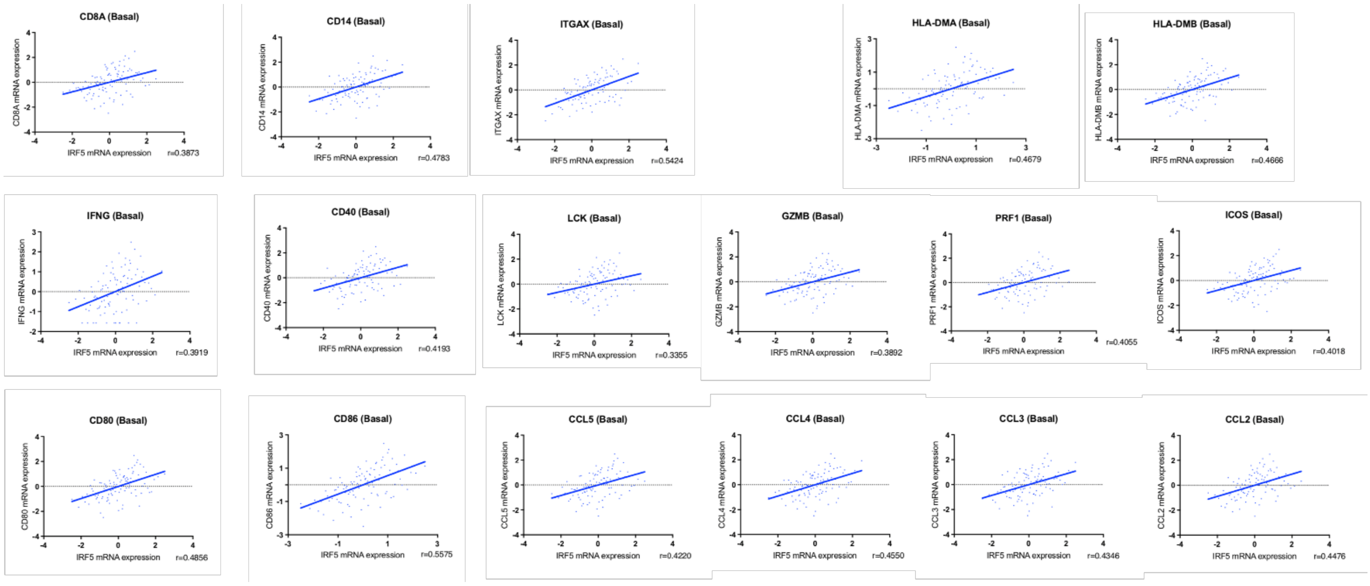


**Supp. Fig. 9.** *IRF5* expression correlation analyses were performed using cBioPortal (TCGA PanCancer Atlas, basal only, n=171) with markers of immune cell activation and function within basal-like breast cancer.

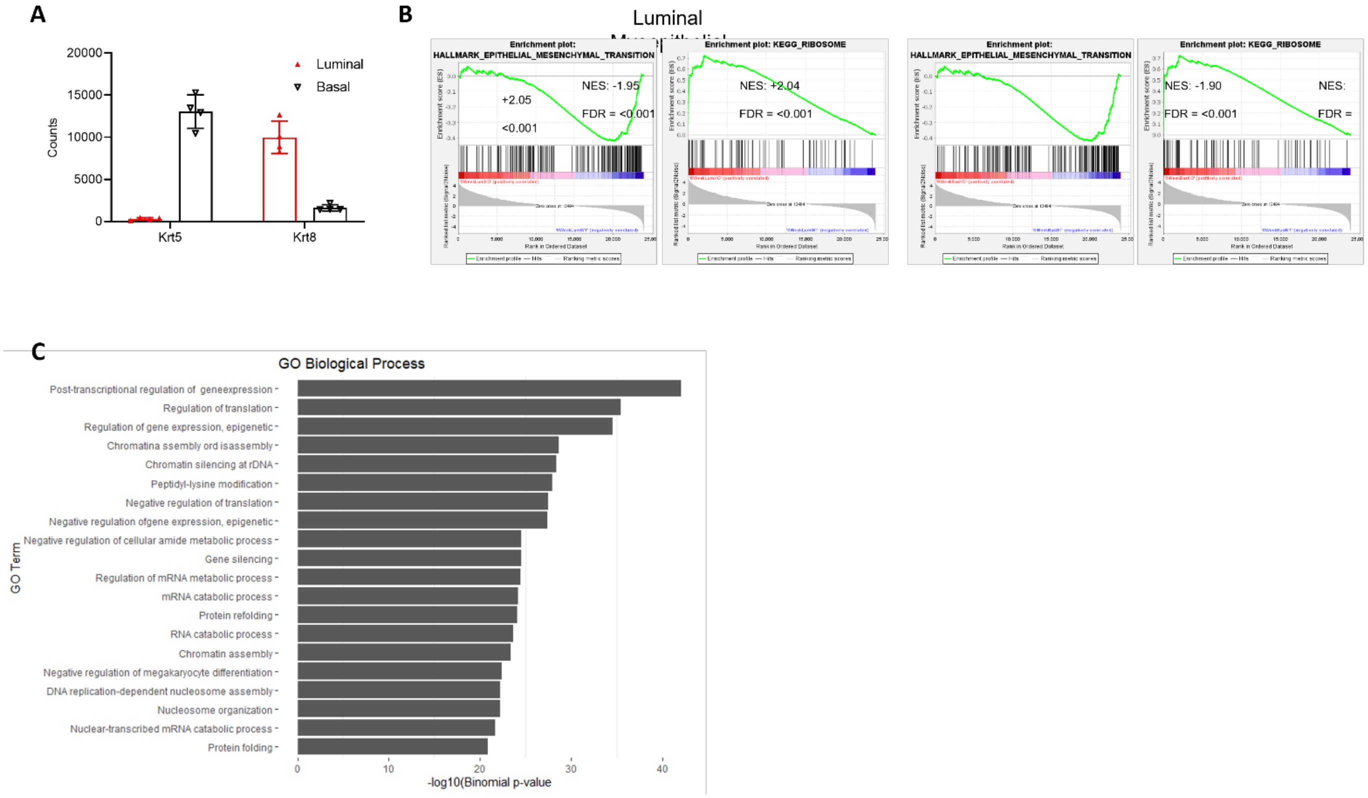


**Supp. Fig. 10.** RNA-seq and ChIP-seq analysis of sorted mammary epithelial cells.

**(A)** Purity of sorted mammary luminal and myoepithelial (basal) cells from 6 weeks-old (n=2/genotype) and 9 months-old (n=2/genotype) WT and *Irf5*$^{-/-}$ mice is shown by RNA-seq analysis of *Krt5* and *Krt8* transcript expression. Ages are plotted together.

**(B)** GSEA of RNA-seq data from sorted luminal and myoepithelial cells of 6 weeks-old WT and *Irf5*$^{-/-}$ mammary glands is shown. Representative terms were chosen with adjusted $p<0.005$. Red is *Irf5*$^{-/-}$ showing positive enrichment for KEGG_Ribosome (luminal, NES +2.04; myoepithelial, NES +2.05); blue is WT showing negative enrichment for HALLMARK_Epithelial to Mesenchymal Transition (luminal, NES -1.95; myoepithelial, NES -1.90). All FDR q-values <0.001.

**(C)** Top significantly enriched gene sets in IRF5 ChIP-seq of mammary epithelial cells identified by gene set enrichment analysis using GO Biological Process.

**Supp. Table I. Tumor incidence in irradiated nulliparous 12 months-old female littermate matched WT and *Irf*5$^{-/-}$ Balb/c mice.**

| | **Pathology** | | | | **Incidence** |
|---|---|---|---|---|---|
| | Normal | ADH | DCIS | IDC | Tumor |
| **γ-Irradiated** | | | | | |
| WT | 8 | 0 | 1 | 0 | 11% |
| *Irf5*$^{-/-}$ | 4 | 0 | 2 | 1 | 43% |

WT, wild-type; ADH, atypical ductal hyperplasia; DCIS, ductal carcinoma in situ; IDC, invasive ductal carcinoma.
